# HUMAN PANCREATIC CANCER CELLS UNDERGO PROFOUND METABOLIC REPROGRAMMING TOWARDS CELLULAR STEMNESS AS ADAPTATION TO INHIBITION OF THE AKT PATHWAY

**DOI:** 10.1101/2020.04.01.020446

**Authors:** Hugo Arasanz, Carlos Hernández, Ana Bocanegra, Luisa Chocarro, Miren Zuazo, Maria Gato, Karina Ausin, Enrique Santamaría, Joaquín Fernández-Irigoyen, Gonzalo Fernandez, Eva Monasterio, Carlos Rodríguez, Idoia Blanco, Ruth Vera, David Escors, Grazyna Kochan

## Abstract

Cancer cells acquire resistance to cytotoxic therapies targeting major survival pathways by adapting their metabolism. The AKT pathway is a major regulator of human pancreatic adenocarcinoma progression. The mechanisms of adaptation to long-term silencing of AKT isoforms of pancreatic cancer cells were studied. Following silencing, cancer cells remained quiescent for long periods of time, after which they recovered proliferative capacities. Adaptation caused profound proteomic changes largely affecting mitochondrial biogenesis, energy metabolism, and acquisition of a number of distinct cancer stem cell (CSC) characteristics depending on the AKT isoform that was silenced. The adaptation to AKT1 silencing drove most de-differentiation and acquisition of stemness through C-MYC down-modulation and NANOG up-regulation, which were required for survival of adapted CSCs. The changes associated to adaptation sensitized cancer cells to inhibitors targeting regulators of oxidative respiration and mitochondrial biogenesis. *In vivo* pharmacological co-inhibition of AKT and mitochondrial metabolism effectively controlled pancreatic adenocarcinoma growth in pre-clinical models.

## INTRODUCTION

Phosphatidyl inositol 3-phosphotase kinase/AKT/molecular target of rapamycin (PI3K/AKT/mTOR) signaling axis constitutes a central pathway involved in progression of pancreatic adenocarcinoma. The main regulators of this pathway are the AKT kinases, which contribute to carcinogenesis, proliferation, migration, angiogenesis and cell survival (Manning and Cantley, 2007). However, AKT-mTOR-targeted therapies are clinically ineffective as tumor cells adapt to overcome inhibition of this key signaling axis (Ben Sahra et al., 2008; El-Khoueiry et al., 2012; Iriana et al., 2016; Javle et al., 2010; Kalender et al., 2010; Kindler et al., 2012; Soares et al., 2013; Wei et al., 2012; Wolpin et al., 2009).

Classically, escape of tumor cells from cytotoxic therapies has been associated to selection of variants with resistance mutations present within a heterogeneous population of tumor cells (Messerschmidt et al., 2017). However, recent evidence indicates that treatment-resistant tumors arise from a population of cancer stem cells (CSC) derived from within a collection of metabolically plastic cancer cells in the tumor (Hermann et al., 2007; Li et al., 2007; Nguyen et al., 2012; Sancho et al., 2015). Indeed, whether treatment-resistant cells come from a pre-existing small population of CSCs, or they are selected by metabolic adaptation through a process of de-differentiation is still a matter of debate. Moreover, therapy-resistant cancer cells can also be selected by other non-genetic mechanisms that include the influence of specific tumor microenvironments through epigenetic regulation (Klemm and Joyce, 2015), distinct mitochondrial content (Guerra et al., 2017; Han et al., 2013; Wallace, 2012) and metabolic reprogramming (Ward and Thompson, 2012) such as the switch from mitochondrial oxidative phosphorylation (OXPHOS) to oxygen-independent glycolysis (*Warburg Effect*) (Vander Heiden et al., 2009). Therefore, it is yet unclear whether inhibition of pro-survival pathways can force cancer cells to directly alter their metabolism through a process of adaptation leading to de-differentiation towards CSCs.

AKT1, AKT2 and AKT3 are three kinase variants with striking similar sequence conservation **(Supplementary Figure 1**). Recent studies indicate that these isoforms do not play redundant roles, but possess specific functions. However, in many instances it is difficult to ascribe the contribution of each isoform to the biology of the cancer cell. While AKT1 is pro-tumorigenic in lung cancer and ErbB2 positive breast cancer (Hollander et al., 2011; Hutchinson et al., 2004; Ju et al., 2007; Linnerth-Petrik et al., 2014; Maroulakou et al., 2007), its silencing is pro-tumorigenic in prostate and other types of breast cancer (Irie et al., 2005; Liu et al., 2006; Virtakoivu et al., 2012; Yoeli-Lerner et al., 2005). In murine breast cancer models, AKT2 abrogation suppresses cell migration, and its expression stimulates motility and invasion in prostate cancer cells *in vitro*, and in breast and ovarian cancer *in vivo*. In contrast, AKT1 and AKT3 have differing effects in these experimental models (Cheng et al., 2007; Irie et al., 2005; Virtakoivu et al., 2012). In murine lung cancer models both AKT2 and AKT3 had anti-proliferative and pro-apoptotic properties (Hollander et al., 2011; Linnerth-Petrik et al., 2014). AKT3 inhibition caused apoptosis and inhibited tumor progression and growth *in vivo* in melanoma and in breast cancer (Chin et al., 2014; Davies et al., 2008).

In pancreatic adenocarcinoma, the consensus is that AKT activities are carcinogenic and pro-tumorigenic. However, there are conflicting results on the exact role of each specific AKT isoform (Altomare and Testa, 2005; Tanno et al., 2001). Here we have studied the mechanism of human pancreatic cancer cells adaptation to sustained inhibition of AKT isoforms, whether this can drive cancer cell de-differentiation towards CSCs and how this process occurs.

## RESULTS

### Adaptation of cancer cells to silencing of AKT isoforms triggers major changes in their proteome

The PI3K-AKT signaling axis is one of the major pathways associated to pancreatic cancer cell progression. However, small molecule inhibitors have demonstrated poor clinical performance. Hence, we decided to investigate the mechanisms by which cancer cells become resistant to its prolonged inhibition by interfering with the expression of each of the AKT isoforms, hoping that this information may also identify the most critical isoform for adaptation. Therefore, cell lines were generated from the human pancreatic ductal adenocarcinoma cell line AsPC-1 in which each AKT kinase isoform was individually silenced. AsPC-1 cells harbor mutations identified in the majority of human adenocarcinomas (K-RAS G35A, p53/p16 inactivation). Lentivectors encoding AKT-targeted shRNAs or shCT (an irrelevant shRNA control) together with puromycin resistance were used to transduce cells and recover puromycin resistant clones. The growth of puromycin-resistant cells was stalled for two months following silencing, after which the cells recovered proliferative capacities. Re-expression of AKT kinases was discarded as an escape mechanism throughout the duration of the study by evaluating silencing of AKT isoforms by western blot at late passages **(Figure 1A)**. Growth kinetics of adapted cell lines were studied by real-time cell monitoring (RTCA). Cell lines adapted to AKT2 and AKT3 silencing still proliferated significantly less than control cells. In contrast, cells adapted to AKT1 silencing recovered higher proliferative capacities than control AKT-non-silenced cells **(Figure 1B)**.

**Figure 1.**
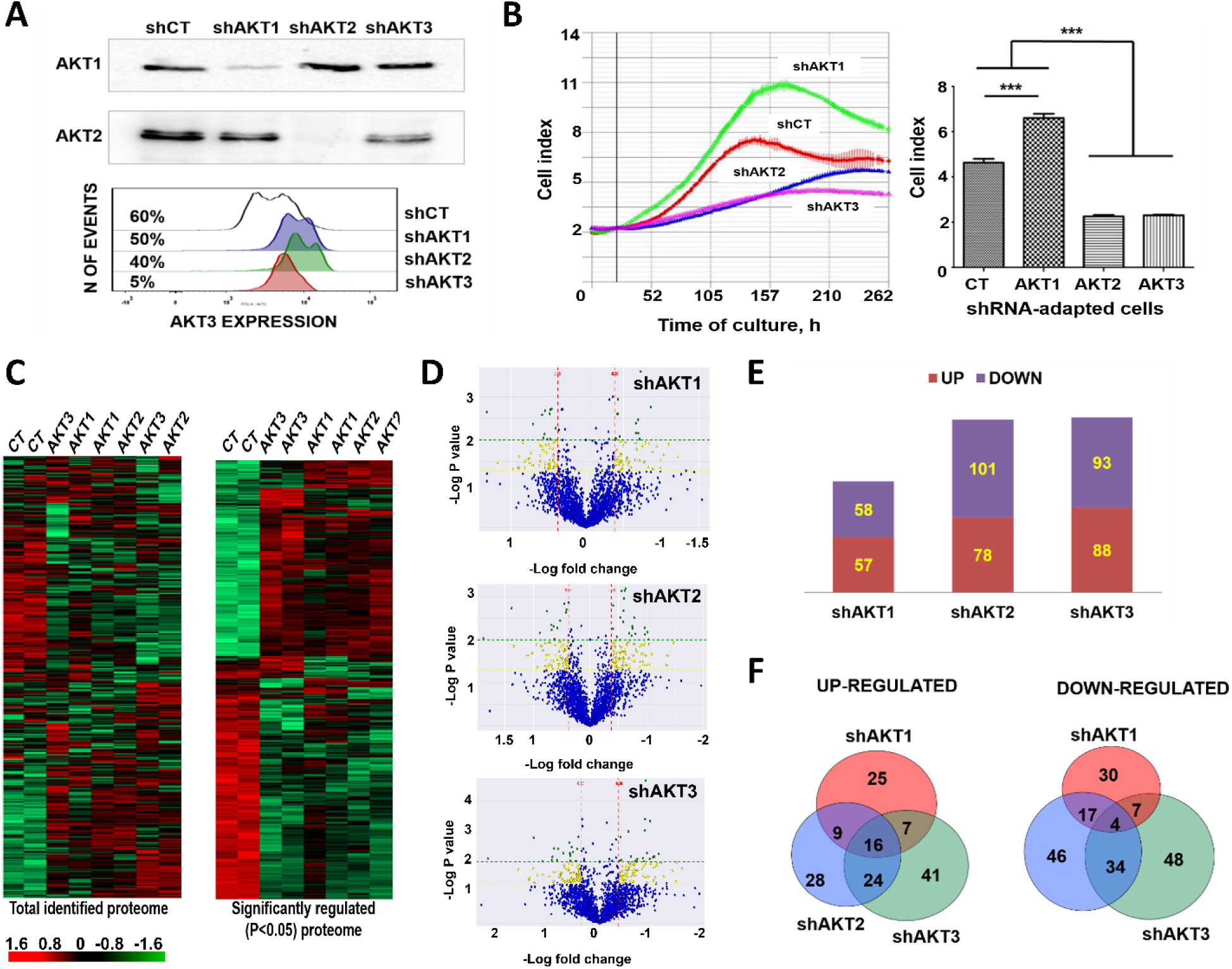
Adaptation of human pancreatic adenocarcinoma cells to specific silencing of AKT isoforms. **(A)** Immunoblots on top, detection of the indicated AKT kinases in cell lines constitutively expressing the shRNA named on top. Below, flow cytometry histogram of AKT3 expression in the cell lines expressing the indicated shRNAs as shown on the right. Percentages of AKT3-expressing cells are shown. Results shown correspond to analyses after more than 20 cell passages after adaptation. **(B)** Left, RTCA graph depicting the growth of the cell lines expressing the indicated shRNAs after adaptation. Right, column graph plotting Cell Index from RTCA data, with means and standard deviations as error bars (4 replicates). Relevant statistical comparisons are shown by ANOVA and Tukey’s pairwise tests. **(C)** Left, cluster analysis of the complete identified proteome from cells adapted to silencing of the indicated AKT isoforms, shown as a heat map. Right, same as left but with significantly regulated (P<0.05) proteins as identified by ANOVA. Red: upregulated genes. Green: downregulated genes. Black: unchanged expression. **(D)** Volcano-plots representing fold-change differences for identified proteins using pair-wise comparisons between shCT-control cells and cell lines adapted to silencing of the indicated AKT isoforms as shown on top of each plot. In blue, proteins without significant changes; Yellow, differentially expressed proteins, P<0.05; Green, differentially expressed proteins, P<0.01. **(E)** bar graph of the number of significantly up-regulated (red) and down-regulated (blue) proteins identified in the volcano plots. **(F)** Venn diagrams of differentially up-regulated (left) or down-regulated (right) proteins in the indicated cell lines with each silenced AKT isoform. Overlaps are shown, together with the number of overlapping identified proteins indicated within the circles. shCT, control shRNA; ***, highly significant (p < 0.001) differences.

To identify the changes that had occurred as a result of adaptation, the proteomes of each cell line were analyzed by quantitative differential proteomics. 3930 proteins were identified with a false discovery rate (FDR) lower than 1%. Cluster analysis of the identified proteomes uncovered major changes that separated cells adapted to silencing of AKT isoforms from control cells expressing an irrelevant shRNA **(Figure 1C)**. Significantly regulated proteins which clustered together the biological replicates were identified for each cell line **(Figure 1C)**. Pair-wise comparisons showed significant quantitative differences between the proteomes **(Figure 1D)**. 115 proteins were differentially regulated in cells adapted to AKT1 silencing compared to control cells (P<0.05), while 179 and 181 were differentially expressed in cells with silenced AKT2 or AKT3, respectively **(Figure 1E)**. Only a minority of down- or up-regulated proteins were shared by cells adapted to silencing of any isoform **(Figure 1F)**. These results indicated that adapted cells had undergone specific proteomic profile changes.

### Adaptation of cancer cells to silencing of AKT isoforms causes profound mitochondrial alterations

Cellular component analyses with differentially regulated proteins for each adapted cell line showed that the most affected organelle was the mitochondrion **(Figure 2A)**. This was confirmed by cluster analysis of the mitochondrion proteome between adapted cells and control cells **(Figure 2B)**. To identify the molecular pathways altered after adaptation compared to control cells, significantly regulated mitochondrial proteins were identified **(Figure 2B)**. Proteins were organized into functional interactomes using STRING and Ingenuity tools, which gave equivalent results. Adaptation to AKT1 silencing increased mTOR regulators and 3 mitochondrial interactomes associated to ATP production, amino acid metabolism, lipid metabolism and mitochondrial DNA replication **(Figure 2C)**. Adaptation to AKT2 and AKT3 silencing activated different interactomes from AKT1-silenced cells, but these were also associated to the regulation of oxidative phosphorylation and mitochondrial protein synthesis **(Figure 2D and 2E)**. Overall, adapted cells had potentiated mitochondrial processes. Increased expression of regulators of mitochondrial DNA replication and protein synthesis was suggestive of mitochondrial biogenesis.

**Figure 2.**
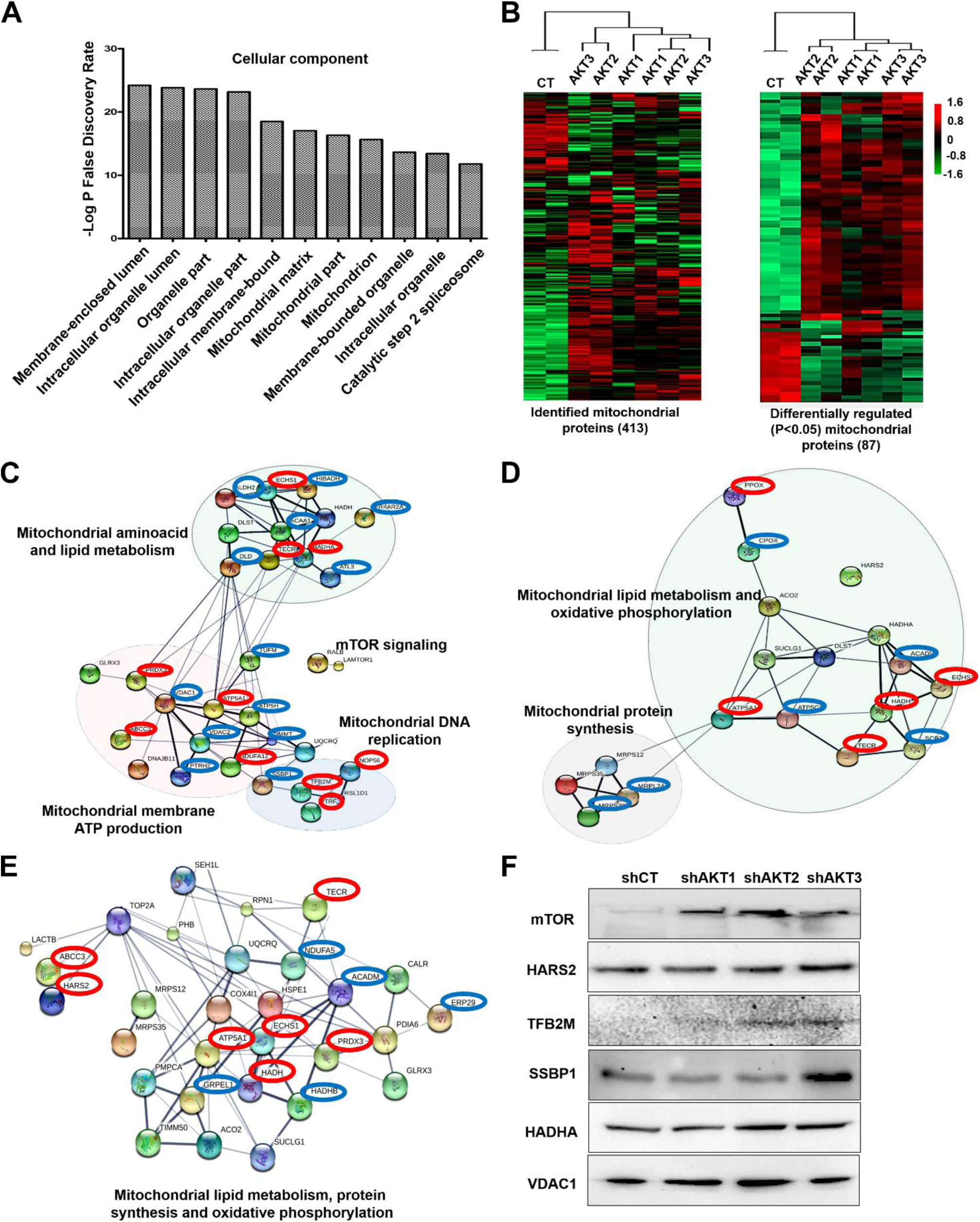
The mitochondrial proteome is significantly altered in cells adapted to AKT silencing. **(A)** Bar graph with the top ten main cellular components in adapted cells to AKT silencing identified using STRING. **(B)** Left, cluster analysis of the identified mitochondrial proteome using two biological replicates per cell line (shown on top), represented by a heatmap. Right, as left but with significantly (P<0.05) regulated mitochondrial proteins between the cell lines (P<0.05) identified by ANOVA. **(C)** Functional interactomes with significantly up-regulated proteins (P<0.05) in cells adapted to AKT1 silencing compared to control cells, and to AKT2 silencing **(D)** and AKT3 silencing **(E)**. In red, common up-regulated proteins present in the three interactomes. In blue, upregulated proteins specific for the adaptation to the indicated AKT kinases. Thin and thick lines shown interactions with medium (0.7) or high (0.9) confidence by STRING. Regulated cellular processes by the functional interactome subgroups are highlighted and indicated. **(F)** Expression of the indicated proteins by western blot in control cells and in cells adapted to silencing of the indicated AKT isoforms.

We tested the regulation of the mTOR pathway and selected protein targets involved in mitochondrial biogenesis **(Figure 2F)**. Samples were loaded in protein gels based on total protein content followed by Ponceau S staining, as standard housekeeping proteins frequently used as loading controls were also differentially regulated in our proteomic data. We confirmed by western blot that mTOR expression was increased in all adapted cells **(Figure 2F)**, as mTOR activation is a known escape mechanism from AKT inhibition (Woo et al., 2017), and a major coordinator of mitochondrial activities, protein synthesis and proliferation (Morita et al., 2015). Overall, differential expression of the mitochondrial proteins TFB2M, SSBP1 HADHA and VDAC1 was generally confirmed **(Figure 2F)**.

To confirm that adaptation caused an increase in mitochondrial mass, MitoTracker Green FM (MG) staining was used as it has been previously demonstrated to be an accurate reporter of mitochondrial mass with negligible phototoxic effects **(Figure 3A)** (Marquez-Jurado et al., 2018; Sancho et al., 2015; Wheaton et al., 2014). Integrated MG intensity in living cells showed a significant increase in mitochondrial mass in adapted cells **(Figure 3B)**. To test if adapted cells relied on mitochondria for growth and survival, we silenced the expression of SSBP1, HARS2 and TFB2M, which regulate mitochondrial DNA replication, protein expression and were upregulated by adaptation to AKT silencing. These targets were silenced with validated shRNAs in AKT1-silenced AsPC-1 cells, as this cell line showed the strongest proliferation and most of the up-regulated interactomes were related with mitochondrial functions. Silencing of these mitochondrial targets caused another proliferative arrest that lasted from 2 to 3 months. Nevertheless, the cells recovered proliferative capacities comparable to the original AsPC-1-shAKT1 cell line **(Figure 3C)** with the exception of cells with silenced TFB2M **(Figure 3C and 3D)**, which had the most significant negative impact over cell growth, with a concomitant decrease of KI67 expression **(Figure 3E)**. Importantly, MG staining confirmed that only TFB2M silencing had significantly reduced mitochondrial content **(Figure 3F)**. Overall, these results corroborated the proteomic data and adapted cells relied on mitochondria for growth.

**Figure 3.**
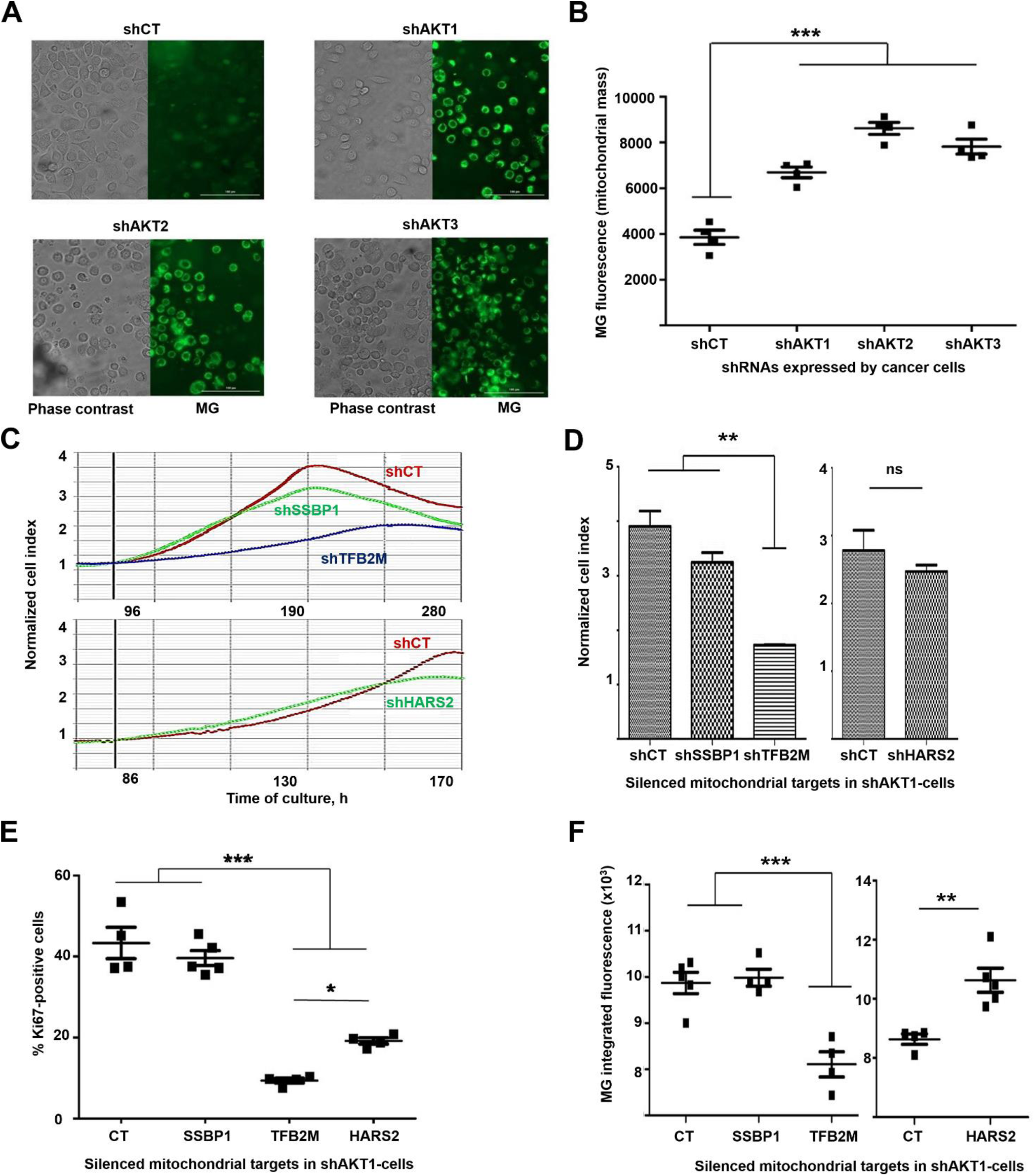
Increased mitochondrial content in cells required for adaptation to AKT silencing. **(A)** Phase contrast and fluorescent microscopy pictures of mitotrack green (MG)-stained cancer cell lines adapted to silencing of the indicated AKT isoforms The control cell line was overexposed to detect background fluorescence signal. Bars within the pictures represent 100 µm. **(B)** Dot plot representing integrated fluorescence intensities for the indicated cell lines, using 4 biological replicates per cell line. Means and standard deviations are also shown. Relevant statistical comparisons are shown by ANOVA and Tukey’s pairwise tests. **(C)** RTCA graphs representing the growth of cells with silenced AKT1 and additional silencing of the indicated mitochondrial proteins. Data are shown with means from duplicate cultures together with error bars. **(D)** Column graphs plotting the Normalized Cell Index from RTCA data corresponding to the left RTCA graphs. Relevant statistical comparisons are shown in the graph by ANOVA and pair-wise comparisons (Tukey’s test). **(E)** Scatter plot of KI-67 expression by flow cytometry for the indicated cell lines. Relevant statistical comparisons are shown within the graph by ANOVA and Tukey’s pairwise tests. **(F)** Scatter plots of integrated MG fluorescence intensity for the indicated cell lines. Data from 5 replicates is shown, with means and error bars (standard deviations). **, ***, indicate very significant (p < 0.01) and highly significant (p < 0.001) differences.

### Adaptation to AKT silencing sensitizes pancreatic cancer cells to mitochondrial disrupting agents

It has been previously shown that cells with a higher mitochondrial content exhibit increased apoptosis, which in turn makes them more susceptible to cytotoxic agents (Akita et al., 2014; Marquez-Jurado et al., 2018). To test whether this was the case for cells adapted to AKT silencing, we quantified the rates of spontaneous apoptosis **(Figure 4A)**. Cells that had adapted to silencing of AKT isoforms showed a slight although significant increase in spontaneous apoptosis compared to control cells. In agreement with this result, cluster analyses of the proteome associated to apoptotic pathways uncovered significant alterations **(Figure 4B)**, with 20 pro- and anti-apoptotic regulators significantly altered. Increased baseline expression of effector caspases 3 and 7 were also found by western blot in adapted cells (not shown). Our results suggested that adapted cells were potentiating mitochondria for survival, which on the other hand could make them more sensitive to apoptosis by mitochondria-disrupting agents. To test whether adapted cells had become sensitized to mitochondria disrupting agents, we evaluated the IC50 of metformin in each cell line by RTCA. Metformin is a potent inhibitor of both the respiratory chain complex I and mTOR (Dowling et al., 2007; Wheaton et al., 2014). The IC50 of metformin was reduced in all cells with silenced AKT isoforms compared to control cells as ascertained by RTCA, and it was significant in AKT1-silenced cells compared to shCT control cells **(Figure 4C**). Tigecycline was then tested as a highly selective inhibitor of the mitochondrial respiratory chain without disrupting the mTOR pathway (Jhas et al., 2013; Skrtic et al., 2011). Cells adapted to AKT1 silencing exhibited the strongest increase in sensitivity to tigecycline (**Figure 4D**), although it did not reach statistical significance **(Figure 4E)**. Overall, these results showed enhanced capacities of mitochondrial-disrupting agents to counteract the growth of cells adapted to silencing of AKT isoforms.

**Figure 4.**
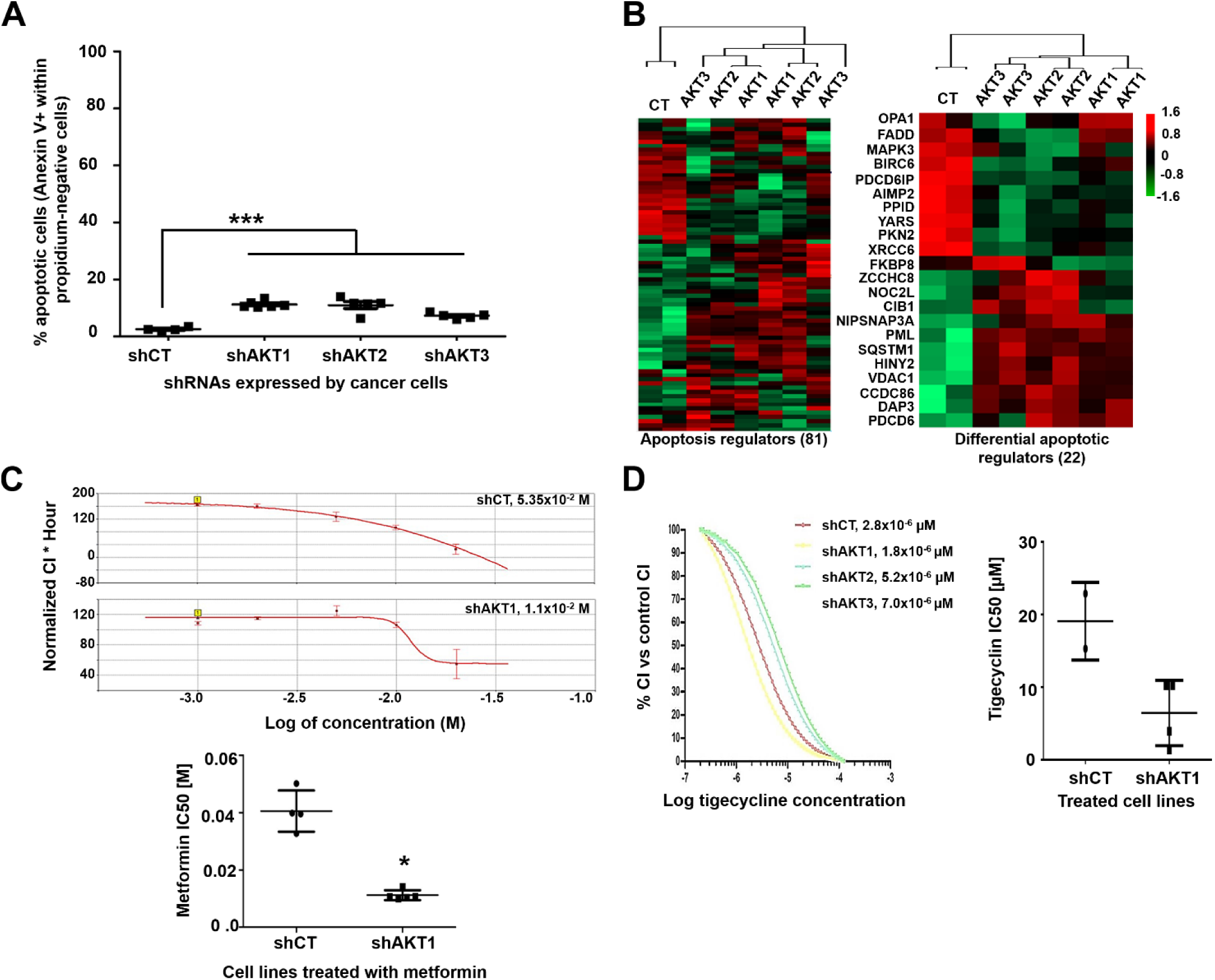
Pancreatic cancer cells with silenced AKT isoforms are sensitive to mitochondria-disrupting agents. **(A)** Dot plot representing the percentage of spontaneous apoptosis in the indicated cell lines. Relevant statistical comparisons are shown by ANOVA and pairwise Tukey’s test. Means and standard deviations are also shown. **(B)** Cluster analysis of proteins involved in regulation of apoptosis (left) and those significantly regulated (right) by silencing of the indicated AKT isoforms. ***, indicate highly significant (p <0.001) differences. **(C)** Top, RTCA IC50 curves of metformin treatments for the indicated cell lines, 4 replicates per concentration. The calculated IC50s for each graph are shown within the graph. Bottom, dot plot graph of IC50 values for metformin treatment calculated from 5 independent experiments as shown on top, in the indicated cell lines. The statistical comparison was performed with Student’s t test. **(D)** Left, RTCA IC50 curves of tigecycline treatments for cell lines with the indicated AKT isoforms silenced, calculated by RTCA. The IC50 values are shown. Right, dot plot graph of IC50 values for tigecycline calculated from 4 experiments. *, ***, indicate significant (P< 0.05) and highly significant (P< 0.001) differences.

### Adaptation to AKT1 silencing causes acquisition of cancer stem cell characteristics

Upregulation of mitochondrial biogenesis and alterations in energy metabolism and mitochondrial respiration by proteomics **(Figures 2 and 3)**, as well as increased sensitivity to metformin and tigecyclin suggested CSCs characteristics in adapted cells To test whether cells had undergone dedifferentiation we first assessed the expression of CD44 and EpCAM, as these represent the two most accepted CSC-associated markers in pancreatic cancer (Heiler et al., 2016; Sancho et al., 2015) **(Figure 5A)**. Interestingly, cells exhibited differential CD44/EpCAM expression profiles specific for the particular AKT isoform that was silenced **(Figure 5B**). Pancreatic cancer cells with silenced AKT1 strongly increased the co-expression of these two CSC markers. AKT3 silencing caused increased expression of CD44 but not EpCAM. Cells adapted to AKT2 silencing upregulated neither CD44 nor EpCAM. These results showed that AKT1-silenced pancreatic cells resembled CSC much closer, and that AKT2-silenced cells were the most differentiated.

**Figure 5.**
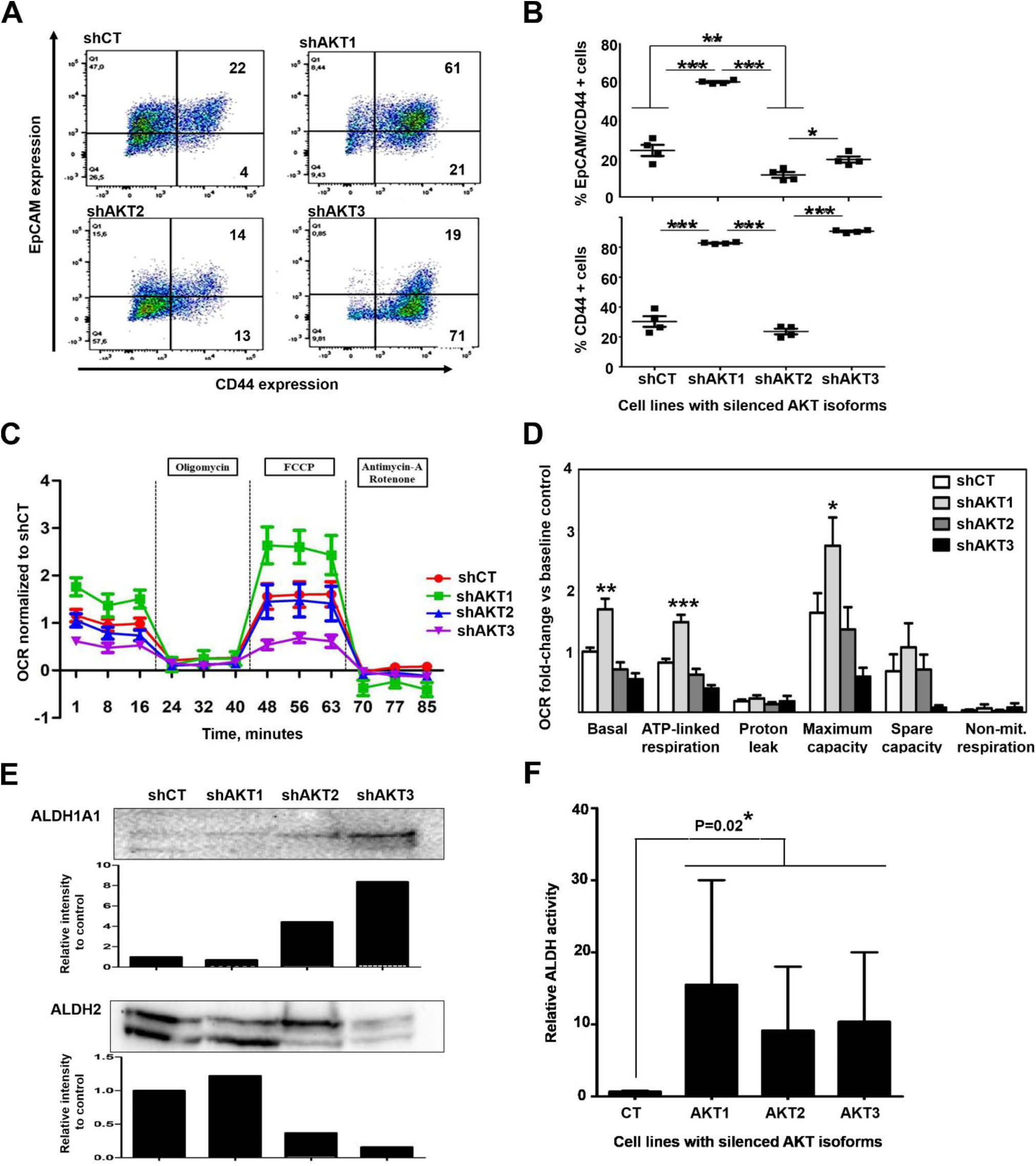
Adapted cells to AKT1 silencing acquire characteristics of cancer stem-like cell. **(A)** Left Flow cytometry density plots of CD44/EpCAM co-expression in the surface of pancreatic cancer cell lines adapted to silencing of the indicated AKT isoform. Percentages of events are shown within each quadrant. **(B)** Scatter plots representing the percentage of cells that co-express CD44/EpCAM (top) or only CD44 (bottom). Data from 4 biological replicates is shown. Relevant statistical comparisons are shown within the graph, calculated by ANOVA and Tukey’s pairwise tests. **(C)** Graph displaying oxygen consumption rates (OCRs) in the indicated cell lines following the addition of oligomycin, FCCP or Antimycin-A/rotenone as shown. Results are shown as means from three technical replicates, together with error bars (SD). **(D)** Bar graph displaying oxygen consumption rate (OCR) for each indicated cell line quantified by SeaHorse XF and normalized to baseline OCR from control cells following the addition of the indicated compounds. Data are shown as means and error bars OCR attributed to each mitochondrial function calculated from (C top), and data shown as means and error bars. **(E)** ALDH1A1 and ALDH2 expression by western blot in the indicated cell lines. Densitometry data on the intensities of both proteins detected by western blot compared to the controls shCT cell line is shown in the bar graphs below each western blot. **(F)** Bar graph representing relative ALDH activity quantified by the Aldefluor flow cytometry staining method, with means and standard deviations as error bars. Relevant statistical comparisons are shown by two-way ANOVA, and Tuckey’s pairwise tests. *, **, ***, indicate significant (P < 0.05), very significant (P<0.01) and highly significant (P < 0.001) differences.

CSCs are highly dependent on oxidative metabolism (Sancho et al., 2015). To study the dependence of AsPC-1 cells with silenced AKT isoforms on oxidative metabolism, real-time oxygen consumption was evaluated by seahorse analyses **(Figures 5C)**. Oxygen consumption by proton efflux, maximal respiration and complete abrogation of mitochondrial respiration were quantified. Only cells with silenced AKT1 exhibited a significant increase in basal and maximal oxygen consumption rates compared to control cells or cells with either AKT2 or AKT3 silenced **(Figure 5D)**.

ALDH activity has been established as a factor responsible for CSC survival, differentiation and resistance to chemo- and radiotherapy. High ALDH1A1 activity has been shown to correlate with CSC phenotype in different cancer types (Tomita et al., 2016). As our proteomic data showed ALDH2 up-regulation in cells adapted to AKT1 silencing compared to the other cell lines including the shCT control, we assessed the expression of both ALDH1A1 and ALDH2 by western blot. Interestingly, ALDH1A1 was overexpressed in cells adapted to AKT3 silencing, while high ALDH2 levels were maintained both in cells adapted to AKT1 silencing and control cells **(Figure 5E)**. Nevertheless, ALDH expression levels do not fully correlate with high enzymatic activity as these enzymes can be inhibited by post-translational modifications such as acetylation (Zhao et al., 2014). Therefore, ALDH activity was quantified in each cell line by flow cytometry using Aldefluor kit **(Figure 5F)**. Interestingly, although there was high variability between experiments, cells adapted to silencing of AKT isoforms showed higher ALDH activities than control cells.

CSCs show changes in basal autophagy as it participates in the acquisition and maintenance of stemness and cell survival in the tumor microenvironment. Autophagy also controls CSC potential for migration and invasion, and promotes resistance to anti-cancer therapies (Nazio et al., 2019; Smith and Macleod, 2019). Pancreatic CSCs activate autophagy to maintain stemness (Valle et al., 2018; Viale et al., 2014). To assess the degree of autophagy in our cell lines, we measured LC3B protein levels by western blot after incubation with the lysosomal inhibitor chloroquine **(Figure 6A)**. LC3B changes from a cytosolic form (LC3B-I) to an autophagosomal membrane-bound form (LC3B-II) during autophagy. Cells with silenced AKT isoforms accumulated more LC3B-II than control cells, indicating that basal autophagy flux was increased in these cells (**Figure 6A and 6B)**. However, the loading β-actin control was found to be regulated to some degree in our proteomic data. Therefore, to directly confirm autophagy activation in living cells, we generated AsPC-1 cells expressing a mCherry/GFP/LC3B (mGLC3B) fusion protein tandem. Using this LC3B version, GFP is less stable in the acidic lysosomal compartment than mCherry, and autophagolysosomes can be easily identified as bright red vesicles by fluorescence microscopy in living cells. Hence, AKT isoforms were silenced in AsPC1-mGLC3B cells, and they were visualized **(Figure 6C)**. Increased number of autophagolysosomes was observed by microscopy especially when AKT1 and AKT2 were silenced. To have a quantitative estimation, we took advantage that in this system autophagy levels proportionally correlate with mCherry/GFP fluorescence ratios (Klionsky et al., 2016). Cells were analyzed by fluorescence microscopy (Cytation 5), and compared to control cells, all cells with silenced AKT isoforms exhibited a higher ratio of mCherry/GFP mean fluorescence intensity, corroborating the increase in autolysosomes (**Figure 6D)**. This increase reached statistical significance for cells with silenced AKT1.

**Figure 6.**
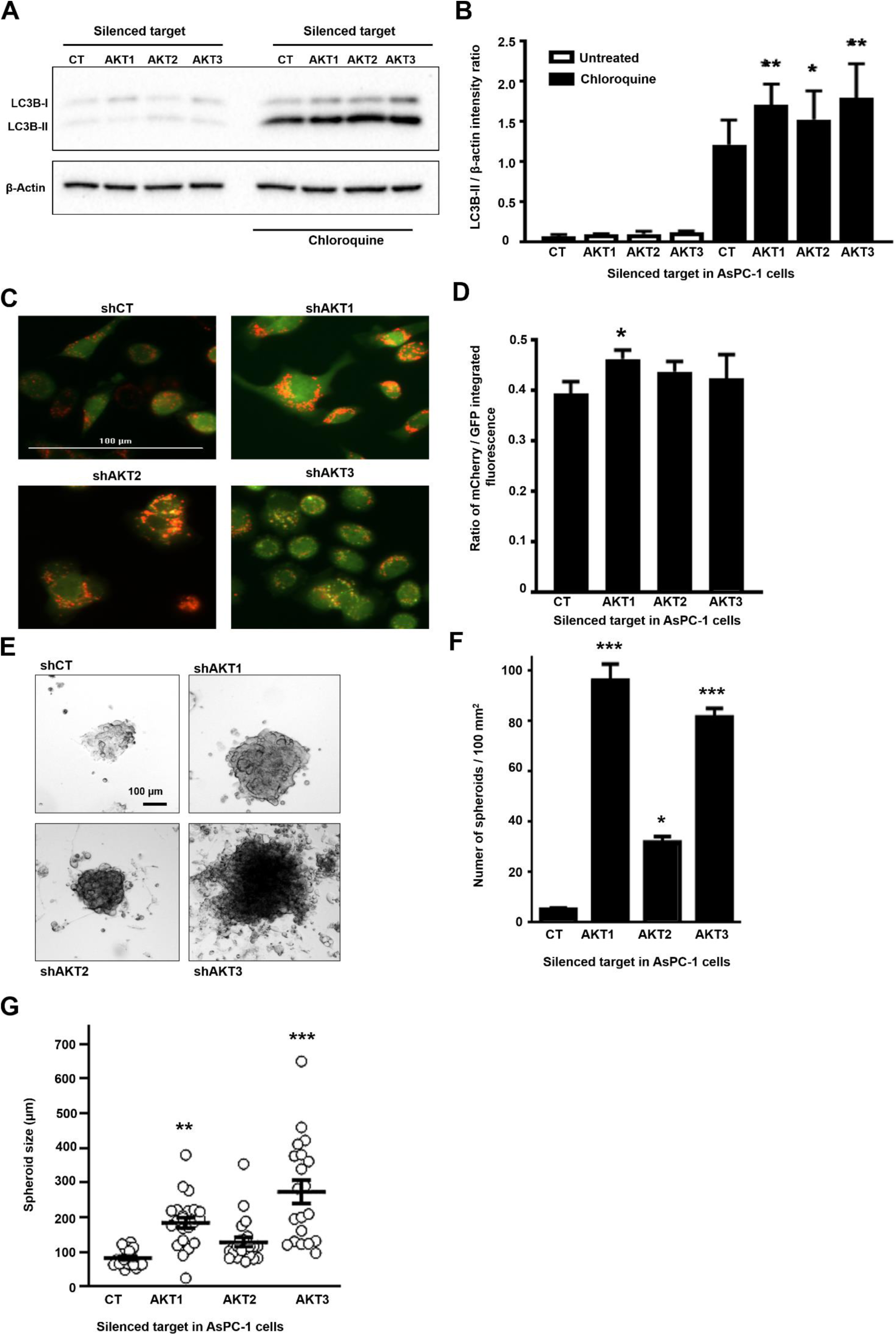
Adaptation of AsPC-1 cells to silencing of AKT isoforms increase autophagy and spheroid growth. **(A)** Western blot with expression of LC3B-I and II forms as indicated in the shown cell lines with or without incubation with chloroquine. **(B)** Bar graph with the ratio of intensities for LC3B-II versus β-actin as determined by densitometry of western blots as in (A). Ratios were obtained from triplicates. Statistical comparisons were performed by ANOVA and pairwise Tukey’s tests between AKT-silenced and control groups. (n=3). **(C)** Fluorescence microscopy of living AsPC-1-mCherry-GFP-LC3B cells with silenced AKT isoforms as indicated, or control cells expressing an irrelevant control shRNA (shCT). Representative 20X pictures with merged GFP and mCherry fluorescence. Autophagolysosomes correspond to bright red vesicles. **(D)** The bar graphs represent the ratio of mCherry/GFP mean fluorescence intensities (n=4). Statistical comparisons were performed by ANOVA and pairwise Tukey’s tests. **(E)** Representative phase contrast pictures of spheroids formed from the indicated AsPC-1 cell lines. **(F)** Bar graph shows the number of spheroids per 100 mm2 obtained from the indicated cell lines. **(G)** Bar graph shows the spheroid size (n=20) from the indicated cell lines. Statistical comparisons were performed by ANOVA and pair-wise Tukey’s test by comparing to control shCT-AsPC-1 cells. *, **, ***, indicate significant (P<0.05), very significant (P<0.01) and highly significant (P<0.001) differences. CT, control shRNA.

One of the most important characteristics of stemness is the increased capacity of cells to grow as spheroids (Noel et al., 2017; Pastrana et al., 2011). Hence, the capacities of the different AsPC-1 cell lines to grow as spheroids were tested. While all the cell lines including shCT-AsPC-1 cells formed spheroids **(Figure 6E)**, cells adapted to silencing of AKT isoforms formed more numerous **(Figure 6F)** and larger spheroids per 100mm^2^ **(Figure 6G)**. Interestingly, the spheroid-forming capacities were significantly higher for AsPC-1 cells with silenced AKT1 or AKT3.

### Adaptation to AKT1 silencing uncovers specific regulation of C-MYC and NANOG

AsPC-1 cells with silenced AKT1 presented most of the features resembling CSC characteristics. To identify potential transcription factors that could regulate the adaptation to AKT1 silencing that could drive the CSC phenotype either by activation or inhibition, we used the TfactS algorithm with differentially regulated proteins as inputs. We used both the whole differential AsPC-1-shAKT1 proteome and also its differential mitochondrial proteome. In both cases, several transcription factors related to cell pluripotency were significantly predicted, but the top significant regulator in both instances was C-MYC **(Figure 7A)**. It has been previously shown that C-MYC activities are down-regulated in pancreatic cancer CSCs (Sancho et al., 2015). Hence, we first assessed C-MYC expression by western blot in AsPC-1 cells with silenced AKT isoforms **(Figure 7B)**. In agreement with a previous report in pancreatic CSCs (Sancho et al., 2015), our results showed lower C-MYC expression also in our adapted cells with silenced AKT isoforms. Hence, to find out if C-MYC functional inhibition was associated to acquisition of a CSC phenotype, constitutively active C-MYCT58A mutant (Thibodeaux et al., 2009) or the dominant negative C-MYCΔHLH mutant (Cartwright et al., 2005) were stably expressed in AsPC-1 cells. Expression of constitutively active C-MYC significantly down-modulated co-expression of CD44/EpCAM **(Figure 7C**) and also exhibited a significant decrease in mitochondrial mass as ascertained by MG staining **(Figure 7C)**. Interestingly, expression of dominant negative C-MYC was not sufficient to drive AsPC-1 de-differentiation. To find out if C-MYC inhibition was a requirement for the undifferentiated state of AsPC-1 cells adapted to AKT1 silencing or their survival, these cells were transduced with a lentivector expressing constitutively active C-MYC T58A together with blasticidin resistance, followed by blasticidin selection. While AsPC-1 control cells could be selected and grown with active C-MYC, it was not possible to select viable AsPC-1 cells adapted to AKT1 silencing with active C-MYC **(Figure 7D)**. These results showed that C-MYC inactivation was absolutely required for survival of pancreatic cancer cells adapted to AKT1 silencing. In agreement with previous studies on pancreatic CSCs (Sancho et al., 2015), our results suggested that C-MYC activities keep AspC1 cells in a differentiated state, but C-MYC inhibition was not sufficient to drive the CSC phenotype although it was necessary for viability of undifferentiated cells.

**Figure 7.**
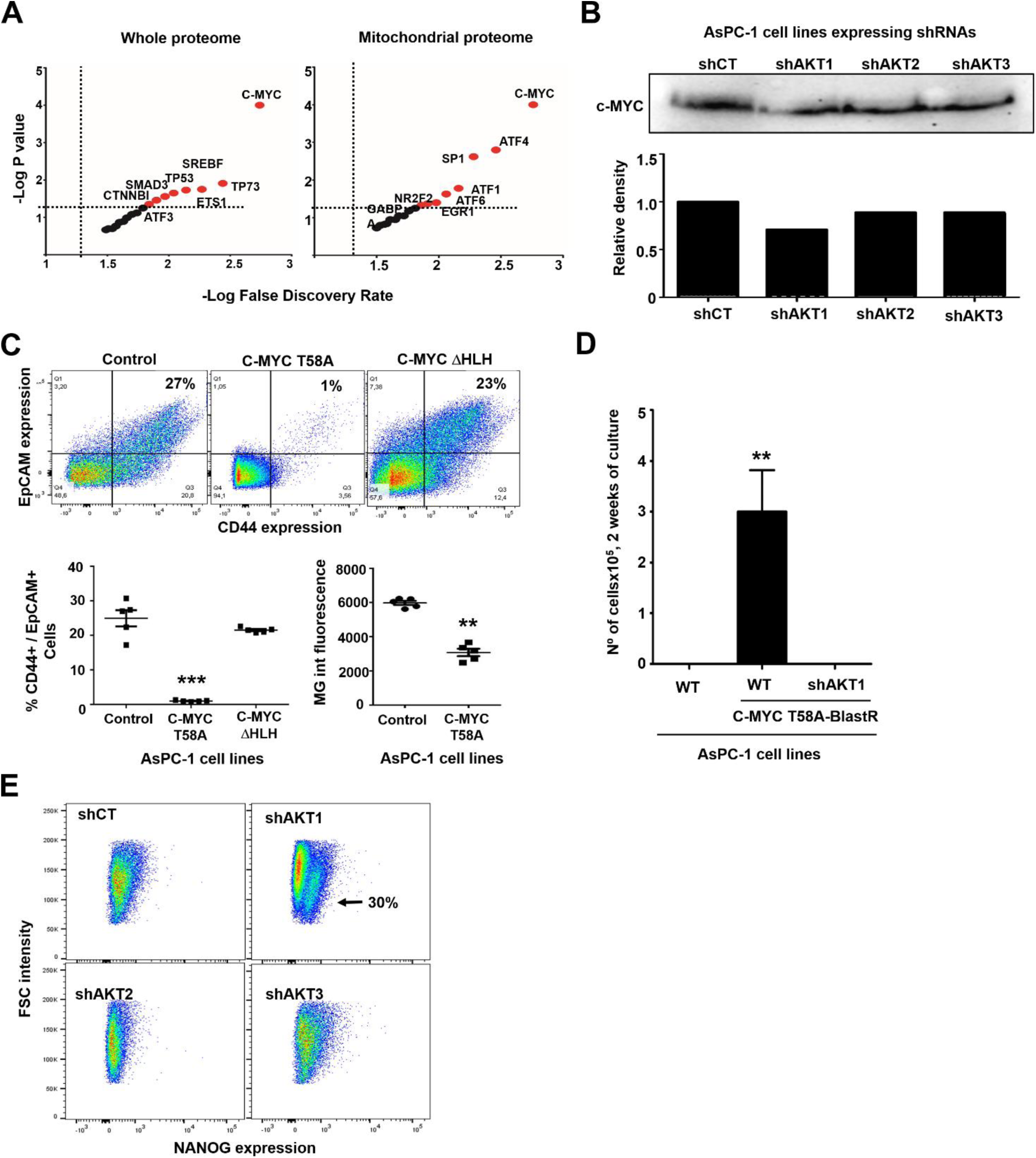
**(A)** Dot plots of transcription factors (indicated within the graph) associated to potential regulation of the proteomes associated to silencing of AKT isoforms compared to shCT-control cancer cells. The left graph represents data from the whole differential proteomes, while the right graph represents data from the mitochondrial differential proteomes. Horizontal and vertical dotted lines separate statistically significant P values and false discovery rates for each identified transcription factor. In red, transcription factors with significant association to the differential proteomes. **(B)** The western blot on top shows C-MYC expression in the indicated AsPC-1 cell lines. The bar graph indicates the band intensity from the western blot above. **(C)** The flow cytometry density plots show CD44/EpCAM expression in the indicated cell lines. Percentages within the graphs show the percentage of CD44-EpCAM double positive AsPC-1 cells; The scatter plot represents the same data from 5 independent experiments. Statistical comparisons were performed by ANOVA and pairwise Tukey’s tests. **(D)** Bar graph representing means and standard deviations of the number of the indicated cell lines transduced with the lentivector expressing active C-MYC and blasticidine resistance, after two weeks of selection. Statistical comparisons were performed by one way ANOVA. **(E)** NANOG expression assessed by flow cytometry in the indicated cells lines. A well-defined NANO-positive population is indicated with an arrow. *, **, ***, indicate significant (p<0.05), very significant (p <0.01) and highly significant (p <0.001) differences, respectively.

Hence, we evaluated the expression of transcription factors that had been previously associated to CSC differentiation, resistance to radio-chemotherapy and disease relapse in various cancers. From a selection of targets, we found that NANOG was specifically expressed in AsPC-1 cells that had adapted to AKT1 silencing **(Figure 7E)**. NANOG has been shown to be responsible for self-renewal and maintainance of pluripotency by transcriptionally repressing genes that drive cell differentiation (Hyslop et al., 2005), and its expression is associated to resistance to conventional therapies (Jeter et al., 2015).

### *In vivo* AKT inhibition together with metformin treatment increases therapeutic efficacy

Our results suggested that dual inhibition of AKT and mitochondrial activities *in vivo* should inhibit tumor progression more effectively than monotherapies, and prolong survival in pre-clinical models of pancreatic cancer. To test this hypothesis, we first subcutaneously transplanted murine pancreatic adenocarcinoma PANC02 cells into C57BL/6 mice, as PANC02 cells harbors K-RAS mutations and exhibit a similar behavior to human pancreatic cancer cells (Partecke et al., 2011). When tumor growth was apparent, mice were subcutaneously administered with suboptimal doses of AKT inhibitor X (10-DEBC hydrochloride, 40mg/1kg) and metformin (40mg/1kg). These doses are well below the lowest published doses in similar experimental models (100 mg/kg, thrice per week for AKT inhibitor, and 100 mg/kg daily for metformin) (Malkomes et al., 2016; Yeo et al., 2018). A group of mice was also treated with their combination (20 mg AKT inhibitor X + 20 mg metformin/1kg), administered twice per week. Mice administered with PBS were used as controls. Only the combo treatment significantly delayed tumor growth **(Figure 8A)**, increased lifespan **(Figure 8B)** and conferred survival advantage **(Figure 8C)**. Treatments with either AKT inhibitor or metformin as monotherapies did not show significant therapeutic efficacies.

**Figure 8.**
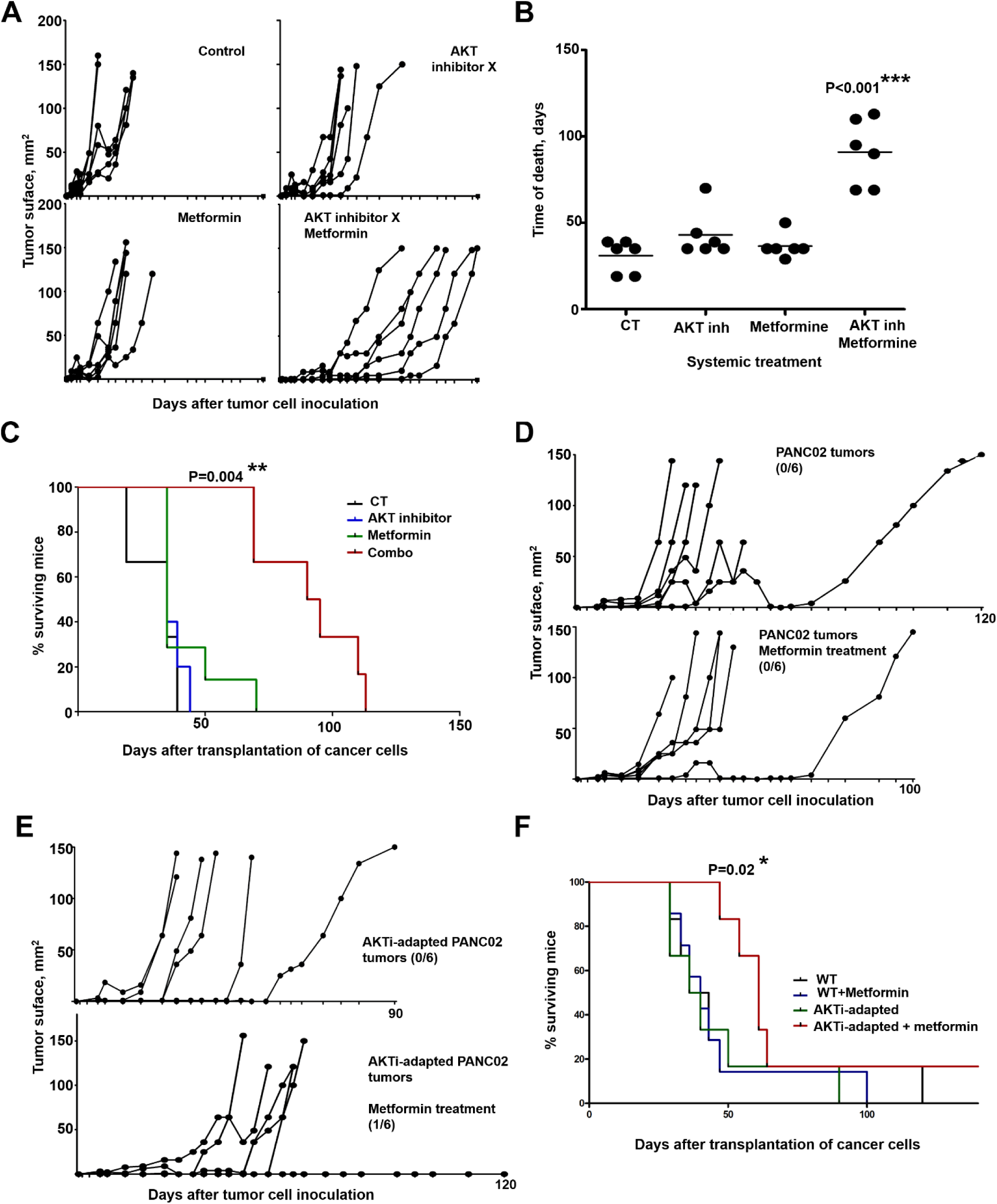
In vivo targeting of AKT and mitochondrial activities synergize in counteracting tumor growth. **(A)** PANC-02 tumor growth in individual mice (n=6) from the indicated treatment groups. Drugs were administered twice subcutaneously. Control group, PBS-treated mice. **(B)** As in (A), but representing the time of death of each mice within the indicated treatment groups. Relevant statistical comparisons were performed with one-way ANOVA and Tukey’s a posteriori test, and shown within the graph. **(C)** Kaplan-Meier survival plot of the treatment groups, as indicated. **(D)** As in (A) but with metformin-only treatment in PANC02 tumors or with AKTi-adapted PANC02 tumors **(E)**. The number of tumor-free mice/total mice are shown within the corresponding treatment groups. **(F)** Kaplan-Meier survival plot of the treatment groups as indicated. Relevant statistical comparisons were performed with.. *, **, ***, indicate significant (p < 0.05), very significant (p < 0.01) and highly significant (p < 0.001) differences.

Then, we adapted PANC02 cells *in vitro* to prolonged inhibition of the AKT pathway by incubation in the presence of progressively increasing concentrations of AKT inhibitor X, until reaching a toxic concentration to non-adapted cells. Similarly to AsPC-1 cells adapted to prolonged AKT1 silencing, AKTi-adapted PANC02 exhibited noticeable phenotypic changes, recovered proliferation capacities comparable to non-adapted cells, and showed a significant increase in mitochondrial mass as assessed by MG staining **(Supplementary figure)**. Groups of mice were then subcutaneously injected with non-adapted **(Figure 8D)** or AKTi-adapted PANC02 cells **(Figure 8E)**, and tumors allowed to grow before starting a suboptimal metformin-only therapeutic regime. Only metformin-treated mice harboring AKTi-adapted PANC02 tumors showed significantly delayed tumor growth **(Figure 8E)**, increased lifespan and survival advantage **(Figure 8F)**. These results confirmed that adaptation to AKT inhibition conferred increased *in vivo* sensitivity to metformin.

## DISCUSSION

Pancreatic adenocarcinoma shows very poor survival after diagnosis, being the fourth most frequent cancer-related death cause. Its treatment is mostly limited to conventional chemotherapy, while targeted therapies exhibit poor clinical outcomes due to progression of resistant cancer cell variants. An increasing number of studies are indicating that the appearance of treatment-resistant cancer cells is dependent on metabolic plasticity and acquisition of CSC characteristics (Hermann et al., 2007; Li et al., 2007; Nguyen et al., 2012; Sancho et al., 2015), apart from the classical selection of cancer cell variants carrying resistance mutations (Messerschmidt et al., 2017). There are also other non-genetic selection mechanisms that can take place through a process of adaptation to specific tumor microenvironments (Klemm and Joyce, 2015; Valle et al., 2018), driven by distinct mitochondrial content (Guerra et al., 2017; Han et al., 2013; Wallace, 2012) or by altering energy metabolism to meet the higher metabolic needs of the tumor cells (Sancho et al., 2015; Vander Heiden et al., 2009).

In the present study we wanted to get insight into the mechanisms of metabolic adaptation of human pancreatic cancer cells to inhibition of AKT, a master regulator of tumor progression and self-renewal of CSCs (Xia and Xu, 2015). However, while most studies use small molecule inhibitors that simultaneously target the three AKT isoforms, we decided to study the implication of each one in the process of molecular adaptation. Instead of genetic ablation of each AKT isoform, these were silenced with RNA interference so as not to completely eliminate their activity. This would resemble more closely the situation of a treated patient with small molecule inhibitors.

The major conclusion from the present study is that human pancreatic cancer cells undergo profound metabolic reprogramming that drives the acquisition of stem cell-like properties at different degrees after adaptation to silencing of AKT isoforms. It is worth noting that silencing of each isoform caused distinct alterations. First, cancer cells remained quiescent for prolonged periods of time after silencing of AKT isoforms. This was expected because AKT directly regulates cell proliferation and survival. After two months, cancer cells regained significant proliferative activities which correlated with a potentiation of mitochondrial functions and mTOR regulators. However, only adaptation to AKT1 silencing caused human pancreatic cancer cells to fully acquire features fulfilling CSC characteristics, including increase in mitochondrial mass, co-upregulation of CD44/EpCAM, increased oxygen consumption, high ALDH activities and autophagy, and enhanced spheroid formation. Apart from these characteristics, lower C-MYC expression and high NANOG expression was confirmed in these cells. Our data agreed with previous reports in which C-MYC activities are down-modulated in pancreatic CSC (Sancho et al., 2015), while Noh and colleagues have shown enrichment of stem-cell phenotypes in human cervical cancer cells by NANOG upregulation (Noh et al., 2012). Moreover, although C-MYC inhibition was not sufficient to drive the CSC phenotype as shown by others, we found that it was an absolute requirement for the viability of CSCs. Cells adapted to AKT3 silencing also showed some CSC characteristics, including CD44 up-regulation, enhanced autophagy and spheroid formation. Therefore, it could be argued that AKT2 is the isoform maintaining de-differentiation towards CSC while AKT1 or ATK3 are silenced. In agreement with this observation, cells adapted to AKT2 silencing do not up-regulate CD44/EpCAM and had the lowest capacities for spheroid growth. These results were not limited to AsPC-1 cells, as murine PANC02 cells adapted to AKT inhibitors also potentiated mitochondrial metabolism to recover proliferative capacities and acquire sensitivity to metformin.

As described before, CSCs are especially sensitive to mitochondria-targeted therapies (De Luca et al., 2015; Lamb et al., 2015a; Lamb et al., 2015b). We found this to be the case with human pancreatic cells adapted to silencing of AKT isoforms, with increased sensitivity towards metformin and tigecyclin and also for mouse PANC02 cells adapted to AKT inhibitor. It is worth noting that tigecycline is an agent shown to selectively target stem cells in CML (Kuntz et al., 2017). Moreover, silencing of TFB2M in cells adapted to AKT1 silencing had a major impact on mitochondrial mass and on its proliferation.

Upregulated ALDH activity was proposed as a CSC marker in different cancers including pancreatic (Ginestier et al., 2007; Marcato et al., 2011; Rasheed et al., 2010). Our proteomic analyses showed that ALDH2 was overexpressed in cells adapted to AKT silencing. It has been recently shown that upregulation of ALDH2 expression in CSCs has a detrimental effect on 5-year survival in different cancer types (Yan and Wu, 2018). Increased autophagy is frequently linked to the CSC phenotype, and it is related with poor prognosis in pancreatic cancer (Fujii et al., 2008). Cells adapted to AKT1 and AKT3 silencing showed increased autophagy, suggesting that AKT2 may promote the CSC phenotype. In agreement with this, cells adapted to AKT1 and AKT3 but not to AKT2 silencing showed enhanced capacities to form spheroids, another characteristic of CSCs.

Our results have significant consequences, and demonstrate that cytotoxic treatments targeting the mTOR/AKT pathway drive de-differentiation towards CSCs after a period of adaptation, without the need of modulating other factors such as a specific tumor environment or acquisition of further mutations. This result strongly suggests that combination therapies targeting the AKT-mTOR signaling axis together with mitochondrial targets may show increased therapeutic capacities *in vivo*. Our preclinical *in vivo* data with PANC02 cells indicates so with a combo of metformin and AKT inhibitor, which significantly delayed tumor growth, increased lifespan and conferred survival advantage. Similarly to our results with human AsPC-1 cells, adaptation of murine PANC02 cells to toxic concentrations of AKT inhibitor led to recovery of proliferation, increased mitochondrial mass and enhanced sensitivity to metformin *in vivo*. Collectively, our data and previous studies support the use of combination therapies that eliminate cancer cells, while inhibiting CSCs arising from adaptation to cytotoxic therapies by targeting their mitochondrial metabolism.

## MATERIALS AND METHODS

### Cell lines

293T were grown as described (Escors et al., 2008). Human pancreatic adenocarcinoma AsPC-1 cells were acquired from American Type Culture Collection (ATCC), and murine pancreatic adenocarcinoma PANC02 cells were a kind gift from Dr. Ignacio Melero. Cells were grown in RPMI supplemented with 10% fetal bovine serum (FBS) and 1% penicillin/streptomycin. Cell growth and survival were monitored in real time using iCELLigence and xCELLigence real-time cell analysis (RTCA ACEA Biosciences) by seeding 20.000 cells and 5.000 cells respectively. Inhibitory concentration 50 (IC50) to metformin and tigecycline for AsPC-1 cells was calculated by RTCA using increasing concentrations as described (Gato-Canas et al., 2015). PANC02 cells were adapted to toxic concentrations of AKT inhibitor X (10-DEBC hydrochloride, SIGMA, CAS number 925681-41-0), by continuous growth with AKTi through a step-by-step increase in concentration until reaching toxic concentrations (50 µM) to non-adapted cells during the course of one month.

Spheroid formation assays were performed as described (Noel et al., 2017; Pastrana et al., 2011). Briefly, cells were trypsinized to obtain single cell suspension, counted, washed with PBS and resuspended in DMEM-F12 medium supplemented with FGF and B27 at density 2×10^3^ cells/ml. Cells were grown for 24 days in nonadherent plates (TC Plate 6 well, suspension F, Sarsted). Afterwards, cells were collected, trypsinised, diluted and new cultures were set. These steps were repeated and cultures analysed for spheroid numbers and sizes.

### Lentivector construction, production and cell transduction

Coding sequences for short hairpin ARNs were designed using Clontech’s bioinformatic tool (http://bioinfo.clontech.com/rnaidesigner/). These sequences were the following: shAKT1: CGCGTGACCATGAACGAGTTTCTCGAGAAACTCGTTCATGGTCACGCGTTTTTG;shAKT2:CGGCTCCTTCATTGGGTACAACTCGAGTTGTACCCAATGAAGGAGCCGTTTTTG;shAKT3:GTAGTCCAACTTCACAAATTGCTCGAGCAATTTGTGAAGTTGGACTACTTTTTTG;shHARS2:ATTAACCCAGCTGCACTATTGTTCAAGAGACAATAGTGCAGCTGGGTTAATTTTTTTG;shSSBP1:GCATGGCACAGAATATCAGTATTTCAAGAG AATACTGATATTCTGTGCCATGTTTTTTG;shTFB2M:GCCCAAAGCGTAGGGAATTATTTTCAAGAGAAATAATTCCCTACGCTTTGGGTTTTTTG. Coding sequences for shRNAs were cloned into the pSIREN lentivector platforms (Gato-Canas et al., 2017; Lanna et al., 2014). The sequence encoding the fusion protein mCherry-GFP-LC3B was synthetized (GeneArt thermofisher) and cloned into pDUAL-Puro and pDUAL-Blast lentivectors (Gato-Canas et al., 2017) under the transcriptional control of the SFFV promoter. Lentivector production and titration were carried out as described elsewhere (Karwacz et al., 2011) (Liechtenstein et al., 2014a) (Selden et al., 2007). Transductions of cell lines were performed with a multiplicity of transduction of 10. Transduced cells were selected with appropriate concentrations of puromycin (Gibco) or blasticidin (Gibco). AsPC-1-mCherry-GFP-LC3B cells were further cloned by limiting dilution.

#### Mitochondrial staining and respiration

Mitochondria were stained with MitoTracker Green FM (Invitrogen) as recommended by the manufacturers. Briefly, cells were incubated at 37°C with pre-warmed MitoTracker staining solution at a concentration of 10 nM for 30 minutes, washed culture medium and fluorescence was quantified by optical microscopy using Cytation™ 5 (Biotek). Mitochondrial respiration was evaluated using Agilent Seahorse XF as described with Agilent Seahorse XF Cell Mito Stress (Divakaruni et al., 2014). ATP-linked respiration was measured in the presence of oligomycin. Maximum respiratory capacity was quantified by the addition of carbonyl cyanide-4 (trifluoromethoxy) phenylhydrazone (FCCP). Non-mitochondrial respiration was measured in the presence of rotenone/antimycin A.

### Autophagy

AsPC1-mGLC3B modified with the indicated shRNAs were grown in 24-well plates for image captures. Representative images of living cells were captured with Cytation 5 fluorescence microscope (20X). Four fields by condition were taken for analyses using filters for mCherry and GFP fluorescence. Cytation 5 software was used to identify cellular events (10 to 100 µm) with GFP fluorescence as reference. Green and red mean fluorescences were quantified for these events and the ratio between mCherry and GFP fluorescence signals was calculated.

#### Immunoblot

Immunoblots were performed as previously described (Lanna et al., 2014). The following antibodies were used: Anti-AKT1 (2H10), Anti-AKT2 (L79B2), anti-AKT3 (L47B1) and anti-mTOR (REF) were purchased from Cell Signaling. Anti-HADHA, anti-VDAC1, anti-HARS2 and anti-SSBP1 were obtained from Abcam. Anti-cMYC was purchased from Invitrogen. Anti-β-actin and anti-LC3B were obtained from Sigma. HRP anti-mouse and anti-rabbit antibodies were purchased from Dako.

#### Flow Cytometry

Surface and intracellular staining were performed using routine protocols as described before (Gato-Canas et al., 2017; Zuazo et al., 2019). The following antibodies were used: Ki67-APC (Biolegend), CD44-APC and CD326/EpCAM-APCVio779 from Miltenyi Biotec. Apoptosis was evaluated by Annexin V / Iodure Propidium staining using the Annexin V-FITC Apoptosis Detection Kit (Invitrogen™ eBioscience™).

ALDH activity was measured by flow cytometry with ALDEFLUOR (StemCell Technologies, Vancouver, BC, Canada) as recommended by the manufacturer. Briefly, 10×10^5^ cells were suspended in Aldefluor assay buffer containing an ALDH substrate (BODIPY-aminoacetaldehyde-diethyl acetate, BAAA-DA) and incubated at 37 °C for 45 min. A negative control for each sample was generated by incubation with the ALDH inhibitor diethylaminobenzaldehyde. Cells were washed with the Aldefluor buffer and analysed by flow cytometry in a flow cytometer (BD Biosciences).

#### Proteomics

Protein extracts derived from adapted cells to silencing of AKT isoforms were homogenized in lysis buffer containing 7 M urea, 2 M thiourea, 4% (w/v) CHAPS, 50 mM DTT. The homogenates were spinned down at 100.000 x g for 1 h at 15°C. Protein concentration was measured in the supernatants with the Bradford assay kit (Biorad).

A shotgun comparative proteomic analysis of cellular proteomes using iTRAQ (isobaric Tags for Relative and Absolute Quantitation) was performed (Unwin et al., 2010). Protein extracts were precipitated with methanol/choloroform, and pellets dissolved in TEAB 0.5M and Urea 6M. Protein quantitation was performed with the Bradford assay kit (Bio-Rad). iTRAQ labeling of each biological condition (in duplicates) was performed according to the manufacturer’s protocol (Sciex). Briefly, equal amounts of cellular proteins (50 μg) were reduced with 50 mM tris (2-carboxyethyl) phosphine (TCEP) at 60 °C for 1 h. Cysteine residues were alkylated with 200 mM methylmethanethiosulfonate (MMTS) at room temperature for 15 min. Protein enzymatic cleavage was carried out with trypsin (Promega; 1:50, w/w) at 37 °C for 16 h. Each tryptic digest was labelled according to the manufacturer’s instructions with one isobaric amine-reactive tags as follows: Tag113, control cells A; Tag114, control cells B; Tag115, shAKT1-A; Tag116, shAKT1-B; Tag117, shAKT2-A; Tag118, shAKT2-B; Tag119, shAKT3-A; Tag121, shAKT3-B. After 1h incubation, labelled samples were pooled and evaporated until < 40 μl in a vacuum centrifuge.

To increase proteome coverage, the peptide pool was fractionated by Pierce Strong Ion Exchange Spin Columns (ThermoFisher). Briefly, the pool mixture was resuspended in 10 mM KH2PO4 20% ACN pH 3 and the column was activated using 10 mM KH2PO4 20% ACN pH 3 0.5 M KCl. After washing steps with 10 mM KH2PO4 20% ACN pH 3, the pool mixture was added to the column and incubated for 5 minutes at room temperature. Different elution solutions were applied (from 10mM to 300mM KCl), collecting 12 peptide fractions. After drying-down the eluted samples, C18 Zip Tip Solid Phase Extraction (Millipore) were perform to purify and concentrated the peptide mixtures. All fractions were reconstituted into 20 μl of 2% acetonitrile, 0.1% formic acid, 98% MilliQ-H20 prior to mass spectrometric analysis.

Peptides mixtures were separated by reverse phase chromatography using an Eksigent nanoLC ultra 2D pump fitted with a 75 μm ID column (Eksigent 0.075 × 250). Samples were first loaded for desalting and concentration into a 0.5 cm length 300 μm ID precolumn packed with the same chemistry as the separating column. Mobile phases were 100% water 0.1% formic acid (FA) (buffer A) and 100% Acetonitrile 0.1% FA (buffer B). Column gradient was developed in a 70 min two step gradient from 2% B to 30% B in 60 min and 30%B to 40% B in 10 min. Column was equilibrated in 95% B for 5 min and 2% B for 15 min. During all process, precolumn was in line with column and flow maintained all along the gradient at 300 nl/min. Eluting peptides from the column were analyzed using an Sciex 5600 Triple-TOF system. Information data acquisition was acquired upon a survey scan performed in a mass range from 350 m/z up to 1250 m/z in a scan time of 250 ms. Top 25 peaks were selected for fragmentation. Minimum accumulation time for MS/MS was set to 75 ms giving a total cycle time of 2.1 s. Product ions were scanned in a mass range from 100 m/z up to 1500 m/z and excluded for further fragmentation during 15 s. The mass spectrometry proteomics data have been deposited to the ProteomeXchange Consortium (http://proteomecentral.proteomexchange.org) via the PRIDE partner repository with the data set identifier PXDXXXXXX. (For reviewers, Username: XXXXXX@XXXXX; Password: XXXXXX. *Data Analysis*. After MS/MS analysis, data files were processed using ProteinPilot™ 5.0 software from Sciex which uses the algorithm Paragon™ (v.4.0.0.0) (Shilov et al., 2007) for database search and Progroup™ for data grouping and searched against Uniprot Human database (September 2016, 70550 entries). The search parameters allowed for cysteine modification by MMTS and biological modifications programmed in the algorithm. Reporter ion intensities were bias corrected for the overlapping isotope contributions from the iTRAQ tags according to the certificate of analysis provided by the reagent manufacturer (Sciex). The peptide and protein selection criteria for relative quantitation were performed as previously described (Liechtenstein et al., 2014b). Several quantitative estimates provided for each protein by ProteinPilot were utilized: the fold change ratios of differential expression between labelled protein extracts; the p-value, representing the probability that the observed ratio is different than 1 by chance. A decoy database search strategy was also used to estimate the false discovery rate (FDR), defined as the percentage of decoy proteins identified against the total protein identification. The FDR was calculated by searching the spectra against the decoy database generated from the target database using a non-lineal fitting method (Tang et al., 2008) and displayed results were those reporting a 1% Global FDR or better. Results were exported for data interpretation. Relative quantification and statistical analysis were provided by the ProteinPilot software, with an additional 1.3-fold change cutoff for all iTRAQ ratios (ratio ≤0.77 or ≥1.3) to classify proteins as up- or down-regulated. Proteins with iTRAQ ratios below the low range (0.77) were considered to be under-expressed, whereas those above the high range (1.3) were considered to be overexpressed.

#### Bioinformatic analyses

Identification of regulatory/metabolic networks was analyzed with STRING (Search Tool for the Retrieval of Interacting Genes) software (http://stringdb.org/) (Szklarczyk et al., 2019). To infer differentially activated/deactivated pathways, proteomic data was further analyzed with QIAGEN’s Ingenuity® Pathway Analysis (IPA) (QIAGEN Redwood City, www.qiagen.com/ingenuity).

#### *In vivo* experiments

C57BL/6 female mice were purchased from The Jackson Laboratory. Approval for animal studies was obtained from the Animal Ethics Committee of the University of Navarra (Pamplona, Navarra, Spain) and from the Government of Navarra. AKT inhibitor X (CAS 925681-41-0) and metformin hydrochloride were purchased from Sigma-Aldrich. Mice were inoculated with PANC02 cells and were treated with suboptimal subcutaneous doses of AKT inhibitor X (1 mg/mouse twice a week), metformin (1 mg/mouse twice a week) or the combination (0.5 mg of each compound/mouse twice a week). PBS-treated mice were used as controls. Tumor size was measured twice a week.

#### Statistics

Proteomic data was analysed with Perseus after normalization, using ANOVA for multicomparisons and t-test for pair-wise comparisons. All other variables under study were tested for normality by Kolmogorov-Smirnov test. Most of the variables were normally distributed with very low variability and homocedasticity, so multi-comparisons were performed by one-way ANOVA followed by pairwise comparisons with Tukey’s tests when required. ALDH activity was compared by two-way ANOVA to eliminate inter-experimental variability. Survival was analysed by Kaplan-Meier plots and Log-Rank tests. Time of death in murine experiments were compared by the non-parametric Kruskall-Wallis test.

---

**Supplementary Figure 1.**
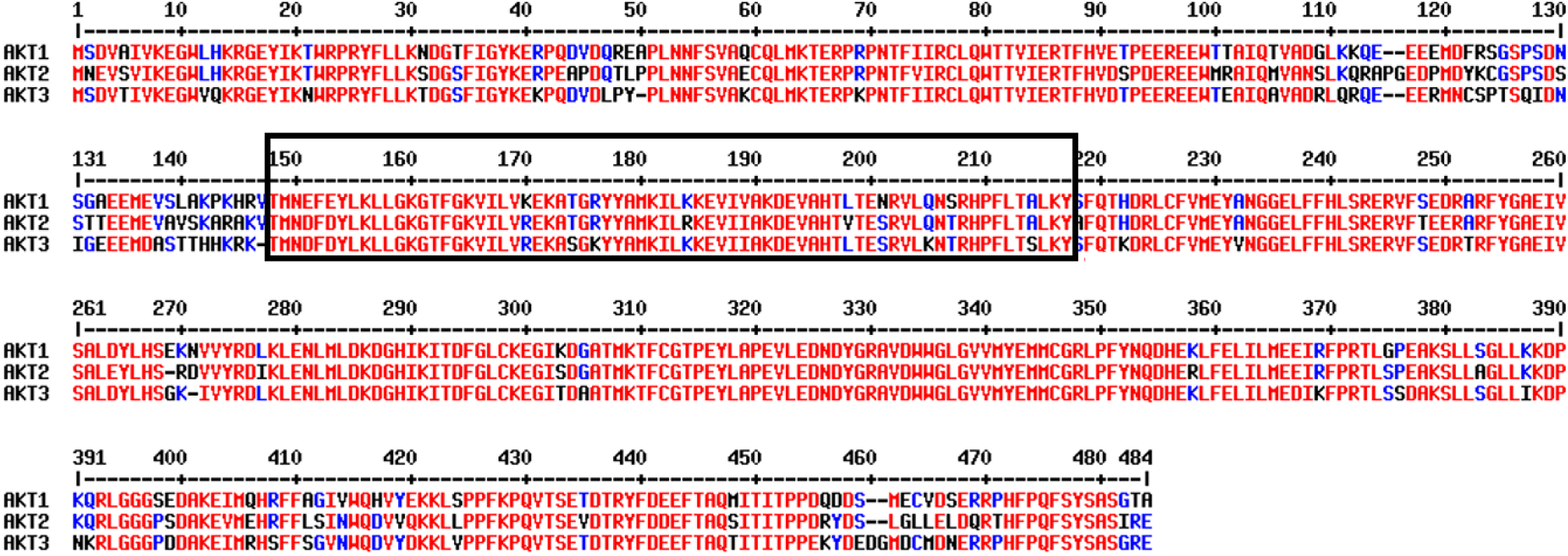
Alignment of aminoacid sequences of the three human AKT isoforms as indicated. Conserved residues are shown in red, and non-conserved residues in blue. The catalytic domain is boxed.

## Acknowledgements

This research was supported by: Asociación Española Contra el Cáncer (AECC, PROYE16001ESCO); Instituto de Salud Carlos III, Spain (FIS project grant PI17/02119); “Precipita” Crowdfunding grant (FECYT); Crowdfunding grant from Sociedad Española de Inmunología (SEI); Government of Navarre project grant in biomedicine (BMED 050-2019). D.E. is funded by a Miguel Servet Fellowship (ISC III, CP12/03114, Spain); H.A. is supported by the Clinico Junior 2019 scholarship from AECC; M.Z. is supported by a scholarship from Universidad Pública de Navarra; and M.G. is supported by a scholarship from the Government of Navarre. L.C.D is supported by a DESCARTHES project grant (Industry department, Government of Navarre).

The authors thank all PRIDE Team for helping with the mass spectrometric data deposit in ProteomeXChange/PRIDE. The Proteomics Unit of Navarrabiomed is a member of Proteored, PRB3-ISCIII, and is supported by grant PT17/0019/0009, of the PE I+D+I 2013-2016 funded by ISCIII and FEDER.

## REFERENCES

Akita, M., Suzuki-Karasaki, M., Fujiwara, K., Nakagawa, C., Soma, M., Yoshida, Y., Ochiai, T., Tokuhashi, Y., and Suzuki-Karasaki, Y. (2014). Mitochondrial division inhibitor-1 induces mitochondrial hyperfusion and sensitizes human cancer cells to TRAIL-induced apoptosis. Int J Oncol 45, 1901–1912.

Altomare, D.A., and Testa, J.R. (2005). Perturbations of the AKT signaling pathway in human cancer. Oncogene 24, 7455–7464.

Ben Sahra, I., Laurent, K., Loubat, A., Giorgetti-Peraldi, S., Colosetti, P., Auberger, P., Tanti, J.F., Le Marchand-Brustel, Y., and Bost, F. (2008). The antidiabetic drug metformin exerts an antitumoral effect in vitro and in vivo through a decrease of cyclin D1 level. Oncogene 27, 3576–3586.

Cartwright, P., McLean, C., Sheppard, A., Rivett, D., Jones, K., and Dalton, S. (2005). LIF/STAT3 controls ES cell self-renewal and pluripotency by a Myc-dependent mechanism. Development 132, 885–896.

Cheng, G.Z., Chan, J., Wang, Q., Zhang, W., Sun, C.D., and Wang, L.H. (2007). Twist transcriptionally up-regulates AKT2 in breast cancer cells leading to increased migration, invasion, and resistance to paclitaxel. Cancer research 67, 1979–1987.

Chin, Y.R., Yoshida, T., Marusyk, A., Beck, A.H., Polyak, K., and Toker, A. (2014). Targeting Akt3 signaling in triple-negative breast cancer. Cancer research 74, 964–973.

Davies, M.A., Stemke-Hale, K., Tellez, C., Calderone, T.L., Deng, W., Prieto, V.G., Lazar, A.J., Gershenwald, J.E., and Mills, G.B. (2008). A novel AKT3 mutation in melanoma tumours and cell lines. British journal of cancer 99, 1265–1268.

De Luca, A., Fiorillo, M., Peiris-Pages, M., Ozsvari, B., Smith, D.L., Sanchez-Alvarez, R., Martinez-Outschoorn, U.E., Cappello, A.R., Pezzi, V., Lisanti, M.P., et al. (2015). Mitochondrial biogenesis is required for the anchorage-independent survival and propagation of stem-like cancer cells. Oncotarget 6, 14777–14795.

Divakaruni, A.S., Rogers, G.W., and Murphy, A.N. (2014). Measuring Mitochondrial Function in Permeabilized Cells Using the Seahorse XF Analyzer or a Clark-Type Oxygen Electrode. Current protocols in toxicology 60, 25 22 21–16.

Dowling, R.J., Zakikhani, M., Fantus, I.G., Pollak, M., and Sonenberg, N. (2007). Metformin inhibits mammalian target of rapamycin-dependent translation initiation in breast cancer cells. Cancer research 67, 10804–10812.

El-Khoueiry, A.B., Ramanathan, R.K., Yang, D.Y., Zhang, W., Shibata, S., Wright, J.J., Gandara, D., and Lenz, H.J. (2012). A randomized phase II of gemcitabine and sorafenib versus sorafenib alone in patients with metastatic pancreatic cancer. Invest New Drugs 30, 1175–1183.

Escors, D., Lopes, L., Lin, R., Hiscott, J., Akira, S., Davis, R.J., and Collins, M.K. (2008). Targeting dendritic cell signalling to regulate the response to immunisation. Blood 111, 3050–3061.

Fujii, S., Mitsunaga, S., Yamazaki, M., Hasebe, T., Ishii, G., Kojima, M., Kinoshita, T., Ueno, T., Esumi, H., and Ochiai, A. (2008). Autophagy is activated in pancreatic cancer cells and correlates with poor patient outcome. Cancer science 99, 1813–1819.

Gato-Canas, M., Martinez de Morentin, X., Blanco-Luquin, I., Fernandez-Irigoyen, J., Zudaire, I., Liechtenstein, T., Arasanz, H., Lozano, T., Casares, N., Chaikuad, A., et al. (2015). A core of kinase-regulated interactomes defines the neoplastic MDSC lineage. Oncotarget 6, 27160–27175.

Gato-Canas, M., Zuazo, M., Arasanz, H., Ibanez-Vea, M., Lorenzo, L., Fernandez-Hinojal, G., Vera, R., Smerdou, C., Martisova, E., Arozarena, I., et al. (2017). PDL1 Signals through Conserved Sequence Motifs to Overcome Interferon-Mediated Cytotoxicity. Cell Rep 20, 1818–1829.

Ginestier, C., Hur, M.H., Charafe-Jauffret, E., Monville, F., Dutcher, J., Brown, M., Jacquemier, J., Viens, P., Kleer, C.G., Liu, S., et al. (2007). ALDH1 is a marker of normal and malignant human mammary stem cells and a predictor of poor clinical outcome. Cell stem cell 1, 555–567.

Guerra, F., Arbini, A.A., and Moro, L. (2017). Mitochondria and cancer chemoresistance. Biochim Biophys Acta Bioenerg 1858, 686–699.

Han, T., Kang, D., Ji, D., Wang, X., Zhan, W., Fu, M., Xin, H.B., and Wang, J.B. (2013). How does cancer cell metabolism affect tumor migration and invasion? Cell Adh Migr 7, 395–403.

Heiler, S., Wang, Z., and Zoller, M. (2016). Pancreatic cancer stem cell markers and exosomes - the incentive push. World J Gastroenterol 22, 5971–6007.

Hermann, P.C., Huber, S.L., Herrler, T., Aicher, A., Ellwart, J.W., Guba, M., Bruns, C.J., and Heeschen, C. (2007). Distinct populations of cancer stem cells determine tumor growth and metastatic activity in human pancreatic cancer. Cell stem cell 1, 313–323.

Hollander, M.C., Maier, C.R., Hobbs, E.A., Ashmore, A.R., Linnoila, R.I., and Dennis, P.A. (2011). Akt1 deletion prevents lung tumorigenesis by mutant K-ras. Oncogene 30, 1812–1821.

Hutchinson, J.N., Jin, J., Cardiff, R.D., Woodgett, J.R., and Muller, W.J. (2004). Activation of Akt-1 (PKB-alpha) can accelerate ErbB-2-mediated mammary tumorigenesis but suppresses tumor invasion. Cancer research 64, 3171–3178.

Hyslop, L., Stojkovic, M., Armstrong, L., Walter, T., Stojkovic, P., Przyborski, S., Herbert, M., Murdoch, A., Strachan, T., and Lako, M. (2005). Downregulation of NANOG induces differentiation of human embryonic stem cells to extraembryonic lineages. Stem cells 23, 1035–1043.

Iriana, S., Ahmed, S., Gong, J., Annamalai, A.A., Tuli, R., and Hendifar, A.E. (2016). Targeting mTOR in Pancreatic Ductal Adenocarcinoma. Front Oncol 6, 99.

Irie, H.Y., Pearline, R.V., Grueneberg, D., Hsia, M., Ravichandran, P., Kothari, N., Natesan, S., and Brugge, J.S. (2005). Distinct roles of Akt1 and Akt2 in regulating cell migration and epithelialmesenchymal transition. J Cell Biol 171, 1023–1034.

Javle, M.M., Shroff, R.T., Xiong, H., Varadhachary, G.A., Fogelman, D., Reddy, S.A., Davis, D., Zhang, Y., Wolff, R.A., and Abbruzzese, J.L. (2010). Inhibition of the mammalian target of rapamycin (mTOR) in advanced pancreatic cancer: results of two phase II studies. BMC Cancer 10, 368.

Jeter, C.R., Yang, T., Wang, J., Chao, H.P., and Tang, D.G. (2015). Concise Review: NANOG in Cancer Stem Cells and Tumor Development: An Update and Outstanding Questions. Stem cells 33, 2381–2390.

Jhas, B., Sriskanthadevan, S., Skrtic, M., Sukhai, M.A., Voisin, V., Jitkova, Y., Gronda, M., Hurren, R., Laister, R.C., Bader, G.D., et al. (2013). Metabolic adaptation to chronic inhibition of mitochondrial protein synthesis in acute myeloid leukemia cells. PloS one 8, e58367.

Ju, X., Katiyar, S., Wang, C., Liu, M., Jiao, X., Li, S., Zhou, J., Turner, J., Lisanti, M.P., Russell, R.G., et al. (2007). Akt1 governs breast cancer progression in vivo. Proceedings of the National Academy of Sciences of the United States of America 104, 7438–7443.

Kalender, A., Selvaraj, A., Kim, S.Y., Gulati, P., Brule, S., Viollet, B., Kemp, B.E., Bardeesy, N., Dennis, P., Schlager, J.J., et al. (2010). Metformin, independent of AMPK, inhibits mTORC1 in a rag GTPase-dependent manner. Cell Metab 11, 390–401.

Karwacz, K., Bricogne, C., Macdonald, D., Arce, F., Bennett, C.L., Collins, M., and Escors, D. (2011). PD-L1 co-stimulation contributes to ligand-induced T cell receptor down-modulation on CD8(+) T cells. EMBO Mol Med 3, 581–592.

Kindler, H.L., Wroblewski, K., Wallace, J.A., Hall, M.J., Locker, G., Nattam, S., Agamah, E., Stadler, W.M., and Vokes, E.E. (2012). Gemcitabine plus sorafenib in patients with advanced pancreatic cancer: a phase II trial of the University of Chicago Phase II Consortium. Invest New Drugs 30, 382–386.

Klemm, F., and Joyce, J.A. (2015). Microenvironmental regulation of therapeutic response in cancer. Trends Cell Biol 25, 198–213.

Klionsky, D.J., Abdelmohsen, K., Abe, A., Abedin, M.J., Abeliovich, H., Acevedo Arozena, A., Adachi, H., Adams, C.M., Adams, P.D., Adeli, K., et al. (2016). Guidelines for the use and interpretation of assays for monitoring autophagy (3rd edition). Autophagy 12, 1–222.

Kuntz, E.M., Baquero, P., Michie, A.M., Dunn, K., Tardito, S., Holyoake, T.L., Helgason, G.V., and Gottlieb, E. (2017). Targeting mitochondrial oxidative phosphorylation eradicates therapy-resistant chronic myeloid leukemia stem cells. Nature medicine 23, 1234–1240.

Lamb, R., Bonuccelli, G., Ozsvari, B., Peiris-Pages, M., Fiorillo, M., Smith, D.L., Bevilacqua, G., Mazzanti, C.M., McDonnell, L.A., Naccarato, A.G., et al. (2015a). Mitochondrial mass, a new metabolic biomarker for stem-like cancer cells: Understanding WNT/FGF-driven anabolic signaling. Oncotarget 6, 30453–30471.

Lamb, R., Ozsvari, B., Lisanti, C.L., Tanowitz, H.B., Howell, A., Martinez-Outschoorn, U.E., Sotgia, F., and Lisanti, M.P. (2015b). Antibiotics that target mitochondria effectively eradicate cancer stem cells, across multiple tumor types: treating cancer like an infectious disease. Oncotarget 6, 4569–4584.

Lanna, A., Henson, S.M., Escors, D., and Akbar, A.N. (2014). The kinase p38 activated by the metabolic regulator AMPK and scaffold TAB1 drives the senescence of human T cells. Nature immunology 15, 965–972.

Li, C., Heidt, D.G., Dalerba, P., Burant, C.F., Zhang, L., Adsay, V., Wicha, M., Clarke, M.F., and Simeone, D.M. (2007). Identification of pancreatic cancer stem cells. Cancer research 67, 1030–1037.

Liechtenstein, T., Perez-Janices, N., Blanco-Luquin, I., Schwarze, J., Dufait, I., Lanna, A., De Ridder, M., Guerrero-Setas, D., Breckpot, K., and Escors, D. (2014a). Anti-melanoma vaccines engineered to simultaneously modulate cytokine priming and silence PD-L1 characterized using ex vivo myeloid-derived suppressor cells as a readout of therapeutic efficacy. Oncoimmunology 3, e29178.

Liechtenstein, T., Perez-Janices, N., Gato, M., Caliendo, F., Kochan, G., Blanco-Luquin, I., Van der Jeught, K., Arce, F., Guerrero-Setas, D., Fernandez-Irigoyen, J., et al. (2014b). A highly efficient tumor-infiltrating MDSC differentiation system for discovery of anti-neoplastic targets, which circumvents the need for tumor establishment in mice. Oncotarget 5, 7843–7857.

Linnerth-Petrik, N.M., Santry, L.A., Petrik, J.J., and Wootton, S.K. (2014). Opposing functions of Akt isoforms in lung tumor initiation and progression. PloS one 9, e94595.

Liu, H., Radisky, D.C., Nelson, C.M., Zhang, H., Fata, J.E., Roth, R.A., and Bissell, M.J. (2006). Mechanism of Akt1 inhibition of breast cancer cell invasion reveals a protumorigenic role for TSC2. Proceedings of the National Academy of Sciences of the United States of America 103, 4134–4139.

Malkomes, P., Lunger, I., Luetticke, A., Oppermann, E., Haetscher, N., Serve, H., Holzer, K., Bechstein, W.O., and Rieger, M.A. (2016). Selective AKT Inhibition by MK-2206 Represses Colorectal Cancer-Initiating Stem Cells. Ann Surg Oncol 23, 2849–2857.

Manning, B.D., and Cantley, L.C. (2007). AKT/PKB signaling: navigating downstream. Cell 129, 1261–1274.

Marcato, P., Dean, C.A., Giacomantonio, C.A., and Lee, P.W. (2011). Aldehyde dehydrogenase: its role as a cancer stem cell marker comes down to the specific isoform. Cell Cycle 10, 1378–1384.

Maroulakou, I.G., Oemler, W., Naber, S.P., and Tsichlis, P.N. (2007). Akt1 ablation inhibits, whereas Akt2 ablation accelerates, the development of mammary adenocarcinomas in mouse mammary tumor virus (MMTV)-ErbB2/neu and MMTV-polyoma middle T transgenic mice. Cancer research 67, 167–177.

Marquez-Jurado, S., Diaz-Colunga, J., das Neves, R.P., Martinez-Lorente, A., Almazan, F., Guantes, R., and Iborra, F.J. (2018). Mitochondrial levels determine variability in cell death by modulating apoptotic gene expression. Nature communications 9, 389.

Messerschmidt, J.L., Bhattacharya, P., and Messerschmidt, G.L. (2017). Cancer Clonal Theory, Immune Escape, and Their Evolving Roles in Cancer Multi-Agent Therapeutics. Curr Oncol Rep 19, 66.

Morita, M., Gravel, S.P., Hulea, L., Larsson, O., Pollak, M., St-Pierre, J., and Topisirovic, I. (2015). mTOR coordinates protein synthesis, mitochondrial activity and proliferation. Cell Cycle 14, 473–480.

Nazio, F., Bordi, M., Cianfanelli, V., Locatelli, F., and Cecconi, F. (2019). Autophagy and cancer stem cells: molecular mechanisms and therapeutic applications. Cell Death Differ 26, 690–702.

Nguyen, L.V., Vanner, R., Dirks, P., and Eaves, C.J. (2012). Cancer stem cells: an evolving concept. Nat Rev Cancer 12, 133–143.

Noel, P., Munoz, R., Rogers, G.W., Neilson, A., Von Hoff, D.D., and Han, H. (2017). Preparation and Metabolic Assay of 3-dimensional Spheroid Co-cultures of Pancreatic Cancer Cells and Fibroblasts. Journal of visualized experiments : JoVE.

Noh, K.H., Kim, B.W., Song, K.H., Cho, H., Lee, Y.H., Kim, J.H., Chung, J.Y., Kim, J.H., Hewitt, S.M., Seong, S.Y., et al. (2012). Nanog signaling in cancer promotes stem-like phenotype and immune evasion. The Journal of clinical investigation 122, 4077–4093.

Partecke, L.I., Sendler, M., Kaeding, A., Weiss, F.U., Mayerle, J., Dummer, A., Nguyen, T.D., Albers, N., Speerforck, S., Lerch, M.M., et al. (2011). A syngeneic orthotopic murine model of pancreatic adenocarcinoma in the C57/BL6 mouse using the Panc02 and 6606PDA cell lines. Eur Surg Res 47, 98–107.

Pastrana, E., Silva-Vargas, V., and Doetsch, F. (2011). Eyes wide open: a critical review of sphere-formation as an assay for stem cells. Cell stem cell 8, 486–498.

Rasheed, Z.A., Yang, J., Wang, Q., Kowalski, J., Freed, I., Murter, C., Hong, S.M., Koorstra, J.B., Rajeshkumar, N.V., He, X., et al. (2010). Prognostic significance of tumorigenic cells with mesenchymal features in pancreatic adenocarcinoma. J Natl Cancer Inst 102, 340–351.

Sancho, P., Burgos-Ramos, E., Tavera, A., Bou Kheir, T., Jagust, P., Schoenhals, M., Barneda, D., Sellers, K., Campos-Olivas, R., Grana, O., et al. (2015). MYC/PGC-1alpha Balance Determines the Metabolic Phenotype and Plasticity of Pancreatic Cancer Stem Cells. Cell Metab 22, 590–605.

Selden, C., Mellor, N., Rees, M., Laurson, J., Kirwan, M., Escors, D., Collins, M., and Hodgson, H. (2007). Growth factors improve gene expression after lentiviral transduction in human adult and fetal hepatocytes. J Gene Med 9, 67–76.

Shilov, I.V., Seymour, S.L., Patel, A.A., Loboda, A., Tang, W.H., Keating, S.P., Hunter, C.L., Nuwaysir, L.M., and Schaeffer, D.A. (2007). The Paragon Algorithm, a next generation search engine that uses sequence temperature values and feature probabilities to identify peptides from tandem mass spectra. Mol Cell Proteomics 6, 1638–1655.

Skrtic, M., Sriskanthadevan, S., Jhas, B., Gebbia, M., Wang, X., Wang, Z., Hurren, R., Jitkova, Y., Gronda, M., Maclean, N., et al. (2011). Inhibition of mitochondrial translation as a therapeutic strategy for human acute myeloid leukemia. Cancer Cell 20, 674–688.

Smith, A.G., and Macleod, K.F. (2019). Autophagy, cancer stem cells and drug resistance. J Pathol 247, 708–718.

Soares, H.P., Ni, Y., Kisfalvi, K., Sinnett-Smith, J., and Rozengurt, E. (2013). Different patterns of Akt and ERK feedback activation in response to rapamycin, active-site mTOR inhibitors and metformin in pancreatic cancer cells. PloS one 8, e57289.

Szklarczyk, D., Gable, A.L., Lyon, D., Junge, A., Wyder, S., Huerta-Cepas, J., Simonovic, M., Doncheva, N.T., Morris, J.H., Bork, P., et al. (2019). STRING v11: protein-protein association networks with increased coverage, supporting functional discovery in genome-wide experimental datasets. Nucleic Acids Res 47, D607–D613.

Tang, W.H., Shilov, I.V., and Seymour, S.L. (2008). Nonlinear fitting method for determining local false discovery rates from decoy database searches. J Proteome Res 7, 3661–3667.

Tanno, S., Tanno, S., Mitsuuchi, Y., Altomare, D.A., Xiao, G.H., and Testa, J.R. (2001). AKT activation up-regulates insulin-like growth factor I receptor expression and promotes invasiveness of human pancreatic cancer cells. Cancer research 61, 589–593.

Thibodeaux, C.A., Liu, X., Disbrow, G.L., Zhang, Y., Rone, J.D., Haddad, B.R., and Schlegel, R. (2009). Immortalization and transformation of human mammary epithelial cells by a tumor-derived Myc mutant. Breast cancer research and treatment 116, 281–294.

Tomita, H., Tanaka, K., Tanaka, T., and Hara, A. (2016). Aldehyde dehydrogenase 1A1 in stem cells and cancer. Oncotarget 7, 11018–11032.

Unwin, R.D., Griffiths, J.R., and Whetton, A.D. (2010). Simultaneous analysis of relative protein expression levels across multiple samples using iTRAQ isobaric tags with 2D nano LC-MS/MS. Nat Protoc 5, 1574–1582.

Valle, S., Martin-Hijano, L., Alcala, S., Alonso-Nocelo, M., and Sainz, B., Jr. (2018). The Ever-Evolving Concept of the Cancer Stem Cell in Pancreatic Cancer. Cancers 10.

Vander Heiden, M.G., Cantley, L.C., and Thompson, C.B. (2009). Understanding the Warburg effect: the metabolic requirements of cell proliferation. Science 324, 1029–1033.

Viale, A., Pettazzoni, P., Lyssiotis, C.A., Ying, H., Sanchez, N., Marchesini, M., Carugo, A., Green, T., Seth, S., Giuliani, V., et al. (2014). Oncogene ablation-resistant pancreatic cancer cells depend on mitochondrial function. Nature 514, 628–632.

Virtakoivu, R., Pellinen, T., Rantala, J.K., Perala, M., and Ivaska, J. (2012). Distinct roles of AKT isoforms in regulating beta1-integrin activity, migration, and invasion in prostate cancer. Mol Biol Cell 23, 3357–3369.

Wallace, D.C. (2012). Mitochondria and cancer. Nat Rev Cancer 12, 685–698.

Ward, P.S., and Thompson, C.B. (2012). Metabolic reprogramming: a cancer hallmark even warburg did not anticipate. Cancer Cell 21, 297–308.

Wei, F., Liu, Y., Bellail, A.C., Olson, J.J., Sun, S.Y., Lu, G., Ding, L., Yuan, C., Wang, G., and Hao, C. (2012). K-Ras mutation-mediated IGF-1-induced feedback ERK activation contributes to the rapalog resistance in pancreatic ductal adenocarcinomas. Cancer letters 322, 58–69.

Wheaton, W.W., Weinberg, S.E., Hamanaka, R.B., Soberanes, S., Sullivan, L.B., Anso, E., Glasauer, A., Dufour, E., Mutlu, G.M., Budigner, G.S., et al. (2014). Metformin inhibits mitochondrial complex I of cancer cells to reduce tumorigenesis. Elife 3, e02242.

Wolpin, B.M., Hezel, A.F., Abrams, T., Blaszkowsky, L.S., Meyerhardt, J.A., Chan, J.A., Enzinger, P.C., Allen, B., Clark, J.W., Ryan, D.P., et al. (2009). Oral mTOR inhibitor everolimus in patients with gemcitabine-refractory metastatic pancreatic cancer. J Clin Oncol 27, 193–198.

Woo, S.U., Sangai, T., Akcakanat, A., Chen, H., Wei, C., and Meric-Bernstam, F. (2017). Vertical inhibition of the PI3K/Akt/mTOR pathway is synergistic in breast cancer. Oncogenesis 6, e385.

Xia, P., and Xu, X.Y. (2015). PI3K/Akt/mTOR signaling pathway in cancer stem cells: from basic research to clinical application. Am J Cancer Res 5, 1602–1609.

Yan, S., and Wu, G. (2018). Could ALDH2(*)2 be the reason for low incidence and mortality of ovarian cancer for East Asia women? Oncotarget 9, 12503–12512.

Yeo, S.K., Paul, R., Haas, M., Wang, C., and Guan, J.L. (2018). Improved efficacy of mitochondrial disrupting agents upon inhibition of autophagy in a mouse model of BRCA1-deficient breast cancer. Autophagy 14, 1214–1225.

Yoeli-Lerner, M., Yiu, G.K., Rabinovitz, I., Erhardt, P., Jauliac, S., and Toker, A. (2005). Akt blocks breast cancer cell motility and invasion through the transcription factor NFAT. Mol Cell 20, 539–550.

Zhao, D., Mo, Y., Li, M.T., Zou, S.W., Cheng, Z.L., Sun, Y.P., Xiong, Y., Guan, K.L., and Lei, Q.Y. (2014). NOTCH-induced aldehyde dehydrogenase 1A1 deacetylation promotes breast cancer stem cells. The Journal of clinical investigation 124, 5453–5465.

Zuazo, M., Arasanz, H., Fernandez-Hinojal, G., Garcia-Granda, M.J., Gato, M., Bocanegra, A., Martinez, M., Hernandez, B., Teijeira, L., Morilla, I., et al. (2019). Functional systemic CD4 immunity is required for clinical responses to PD-L1/PD-1 blockade therapy. EMBO Mol Med 11, e10293.

